# Concentration and dosage sensitivity of proteins driving liquid-liquid phase separation

**DOI:** 10.1101/2021.02.19.430946

**Authors:** Nazanin Farahi, Tamas Lazar, Shoshana J. Wodak, Peter Tompa, Rita Pancsa

## Abstract

Liquid-liquid phase separation (LLPS) is a molecular process that leads to the formation of membraneless organelles (MLOs), i.e. functionally specialized liquid-like cellular condensates formed by proteins and nucleic acids. Integration of data on LLPS-associated proteins from dedicated databases revealed only modest overlap between them and resulted in a confident set of 89 human LLPS driver proteins. Since LLPS is highly concentration-sensitive, the underlying experiments are often criticized for applying higher-than-physiological protein concentrations. To clarify this issue, we performed a *naive* comparison of *in vitro* applied and quantitative proteomics-derived protein concentrations and discuss a number of considerations that rationalize the choice of apparently high *in vitro* concentrations in most LLPS studies. The validity of *in vitro* LLPS experiments is further supported by *in vivo* phase-separation experiments and by the observation that the corresponding genes show a strong propensity for dosage sensitivity. This observation implies that the availability of the respective proteins is tightly regulated in cells to avoid erroneous condensate formation. In all, we propose that although local protein concentrations are practically impossible to determine in cells, proteomics-derived cellular concentrations should rather be considered as lower limits of protein concentrations, than strict upper bounds, to be respected by *in vitro* experiments.

## 1. Introduction

An important recent discovery in the field of molecular cell biology is that the formation of biomolecular condensates in living cells is driven by a reversible process called liquid-liquid phase separation (LLPS) [1]. These condensates, the so-called membraneless organelles (MLOs), represent distinct liquid phases selectively enriched in certain macromolecules and fulfil essential cellular functions under normal conditions and in response to stress [2–5]. Nucleoli, stress granules, P-bodies, germ granules, postsynaptic densities, heterochromatin and many other long-known or recently discovered cellular compartments that belong to this category [6] have been reported in organisms from all kingdoms of life [7,8]. The functional benefits of MLOs do not directly derive from the individual role of their constituent molecules, but emerge from their collective behavior [4]. Therefore, this recently recognized process is now considered as a fundamental mechanism employed by living cells to cost-efficiently organize and reorganize cellular space and material according to functional needs [5].

A number of studies have demonstrated that MLOs exhibit fluid-like dynamics and general behavior [9–11]. In contrast to classical organelles, MLOs are reversible supramolecular assemblies that enable thermodynamically driven exchange of material with the surrounding solvent [12]. It has also been observed that, in perturbed cellular states, phase-separated liquid-like structures can transition into less dynamic hydrogels or solid-like protein aggregates. The latter often contain long filaments resembling amyloid fibers, involved in many neurodegenerative diseases [13] such as amyotrophic lateral sclerosis [14,15], frontotemporal dementia [16] and Alzheimer’s disease [17–19], drawing attention to the potential pathological roles of liquid condensates.

LLPS is a complex and ill-understood process driven by multivalent weak interactions [12]. It has a very heterogeneous molecular background, both in terms of the interacting macromolecular modules and the contributing molecular driving forces [2,3,20,21]. In addition, protein-protein and protein-nucleic acid interactions can both play a role in scaffolding the condensates. Protein-protein assemblies are driven by interactions between diverse protein modules, such as homooligomerization by domains or coiled coil regions, interactions between intrinsically disordered regions (IDRs), domain-domain, domain-motif, or PTM-controlled molecular interactions [21]. The associated protein-nucleic acid interactions are also diverse, with RNA/DNA length- and sequence-specific [22], as well as structure-specific [22,23] interactions playing a role, mediated by well-folded domains (such as RNA-recognition motifs or KH domains) [24] or IDRs (such as RGG boxes [25,26]) of the participating proteins. At the atomic level, electrostatics, hydrophobic interactions, cation-π or π-π interactions, or a combination of those, are reported to play key roles [2,3,20,21,27,28].

The ability to undergo LLPS may be a property of many macromolecules under specific conditions, many of which may not be encountered in living cells. Therefore, only a subset of proteins will phase separate to form MLOs under physiologically relevant conditions [3]. Importantly, LLPS is particularly sensitive to environmental conditions, such as ionic strength, temperature, pH, crowding or the concentrations of the participating macromolecules [3,29]. Therefore, the reversible formation of condensates often acts as an ultrasensitive mechanism for sensing subtle changes in the intracellular milieu [30] or as a buffering process enabling the maintenance of fine-tuned intracellular concentrations of their constituent macromolecules [3]. Their sensitivity of LLPS enables sophisticated regulatory mechanisms via cellular parameters, such as pH, temperature, ATP or ion concentrations. Regulatory mechanisms controlling the availability of competitive binders, PTMs or alternatively spliced forms of the condensate components are likewise common [29,31–33].

To categorize a protein as “phase separating” therefore requires a system-level understanding of the phase diagram of the process in the cell, and the influence of cellular parameters and states thereof. But such analyses remain extremely challenging because relevant key parameters are either not known or cannot be controlled. Instead, researchers turn to investigating LLPS in the test tube, where conditions can be readily controlled. There is, however, no guarantee that the findings of *in vitro* experiments accurately represent the process in living cells, where additional molecular species may be present and various regulatory mechanisms may be at play. It is therefore crucial that *in vitro* observations on condensate formation be confirmed by suitable *in vivo* experiments.

The flurry of publications reporting on new, experimentally verified cases of LLPS, and the mounting interest in the LLPS process created the motivation to develop dedicated databases. These include databases such as NsortDB [34], MSGP [35] and RNA granule database [36] that are limited to certain species or MLOs and include proteins based solely on evidence of localization to those MLOs, irrespective of their potential role in LLPS. Of much wider scope are four LLPS-dedicated databases: PhaSePro [37], LLPSDB [38], DrLLPS [39] and PhaSepDB [40], reported in the NAR database issue of 2020. These databases curate data from the literature and aim to provide rich annotations on the LLPS process from all studied species and MLOs. Most of them also annotate the role(s) of proteins involved, thereby offering the scientific community data that can be integrated with information on protein sequence, 3D structures and functional annotations, and used in various bioinformatics analyses. While the general focus of the four databases and the types of published studies that they curated are very similar, the data they store and the annotations they provide differ substantially (see [41,42] for their detailed comparison). This is not too surprising considering the inherent complexity of LLPS in cells and the ensuing challenges of extracting meaningful and consistent information from the literature on the underlying molecular players and conditions.

To better understand what causes these noted differences, we first established critical definitions for the four major LLPS-related protein categories (LLPS driver, co-driver, regulator and client), and outlined the experimental approaches typically used in LLPS studies. Through these, we highlighted the main issues that may confound consistent interpretation of the results. With these issues in mind, we then scrutinized data in these wider-scope LLPS databases by, among other things, attempting to consolidate their annotated protein entries. We uncovered reasons of limited overlap between the annotated proteins and inconsistencies in database annotations, partially due to contradictions among roles assigned to proteins in LLPS. A closer analysis of annotations allowed us to resolve these contradictions and assemble a confident dataset of 89 human LLPS driver proteins whose central role in LLPS is sufficiently supported by physiologically relevant experiments.

Based on the exceptional sensitivity of LLPS to protein concentrations, of potential concern could be the apparent systematic discrepancy we observe between protein abundance in cells and the range of protein concentrations used in *in vitro* LLPS experiments. However, we highlight several considerations that help close this gap and apparently relieve these concerns. To provide direct evidence to these points, we show that genes of the described LLPS drivers are extremely dosage sensitive, i.e. their (local) cellular concentrations are fine-tuned to an optimal range for LLPS, deviations from which induce possibly deleterious phase transitions. In conclusion, we suggest that our integrated, filtered dataset of human LLPS proteins will inspire further large-scale LLPS studies, and our results highlight important factors that need to be taken into account when designing, interpreting or judging the biological relevance of LLPS experiments.

## 2. Results and Discussion

### 2.1 Interpretation of LLPS experiments to define the roles of proteins in the formation and integrity of MLOs

Due to the heterogeneity of the underlying driving forces, molecular mechanisms and macromolecules contributing to LLPS, comprehensively elucidating functional-regulatory details of LLPS systems is a highly challenging task. Experimental approaches to phase-separating systems have been thoroughly reviewed recently [3,11], here we rather focus on the interpretation of the results with regards to the functional roles of LLPS-related proteins, as outlined below.

In principle, a broad range of techniques has been applied and adapted for detecting and/or characterizing LLPS. As our purpose here is to clarify distinct roles of phase-separating proteins and understand the reasons of substantial differences between data in the four LLPS-related databases, we first clarify some cornerstone concepts that are not always articulated in LLPS studies.

1. Strictly speaking, the capacity to phase separate is not a binary classifier, i.e., not the intrinsic property, of a protein, rather a contextual property of the protein and its environment (temperature, pH, partners... etc). For considering a protein “phase-separating”, this behavior has to be approached under “native” or “native-like” conditions that incorporate all important constituents.
2. Proteins have distinct roles (types of contributions) in phase separation, which may even differ between conditions in the test tube and the cell. Roughly, we may distinguish drivers (scaffolds), co-drivers (co-scaffolds), regulators and clients, as defined below. There are various different experimental approaches that contribute to identifying the role of a given protein in LLPS.
3. Phase separation depends on the concentration of the protein (among other parameters), i.e. practically any protein can be made to phase separate at sufficiently high concentrations if the environmental conditions allow it. From the biological (i.e., not polymer-chemical) point of view, we should accept a protein as phase separating if it does phase separate at concentrations compatible with physiological (or pathological) conditions.
4. LLPS is not the equivalent of biomolecular condensation in general. Whereas LLPS is the process of demixing that leads to the formation of dense liquid droplets, condensation is a much broader category that encompasses all reactions of physical assembly, also including gelation, crystallization, clustering, polymerization and amorphous or amyloid aggregation.

As a rule of thumb, we may state that for a complete and correct classification of a given protein with regards to the role (if any) it plays in LLPS, an integration of multiple experimental approaches need to be achieved. We should appreciate that each approach provides different and often complementary information, i.e. in a sense they all have “advantages” and “disadvantages”. In general, the major advantage of *in vitro* experiments is that the components of the system are known and they can be perfectly controlled, whereas their disadvantage is that conditions are over-simplified and cannot exactly recapitulate physiological conditions (in terms of partners, post-translational modifications, metabolites, cellular crowding, etc...). On the other hand, the major advantage of an *in vivo* measurement is that it does report on the LLPS behavior under truly physiological conditions (unless protein(s) are severely overexpressed), enabling it to address the biological relevance of LLPS. Their major disadvantage is that due to the underlying and largely hidden cellular complexity and lack of control on many parameters, many details (in terms of the necessity, role and influence of components) cannot be ascertained.

In general, LLPS systems can only be sufficiently explored, the underlying molecular mechanisms fully uncovered and the roles of the components precisely determined, if *in vivo* and *in vitro* experiments are used in combination and the liquid material state of the resulting condensates is verified. To this end, we first define the distinct roles proteins play in the process of LLPS by defining the major LLPS-related protein categories. For each of them, we provide a short “operational” description of what experimental evidence is required to ascertain them, which then enables us to position the experimental approaches with regards to the type and interpretation of information they provide.

#### Driver (scaffold)

is a protein that can phase separate on its own (given appropriate, native-like conditions) to a dense liquid droplet, without the need for other macromolecular partner(s). We consider “driver” and “scaffold” synonymous. If the presence of RNA is mandatory for LLPS, we consider the protein and RNA “co-drivers” (see next section).

*In vitro*, a driver is observed to phase separate when conditions (such as temperature, concentration, PTM state, the presence of a crowder... etc.) are right. In this framework, small molecules facilitating the LLPS of a driver (e.g. particular buffer, salt, metabolite) are considered “conditions”, and not co-drivers of LLPS. *In vivo*, overexpression of a driver, its oligomerization (driven by a PTM or optogenetics) or translocation to another organelle is sufficient to cause LLPS, i.e. the appearance of cellular puncta (membraneless organelles). Physiological cellular/local concentrations are preferable, and disappearance of the organelle upon deletion/silencing of the protein is not necessarily conclusive.

A ***co-driver*** is a macromolecule (protein, RNA or DNA) that strictly requires another co-driver for phase separation. The two (or more) co-drivers are usually equally mandatory for LLPS (e.g. partners in signalosomes built on domain-motif interactions), however, in some cases one co-driver can phase separate on its own in very high concentrations and its partner is required to lower its saturation concentration to physiologically relevant levels (e.g. RNA in some ribonucleoprotein droplets). A co-driver has a different role from a “regulator”, which also promotes the LLPS of its partner, but it does not physically take part in LLPS, i.e. will not be part of the resulting MLO scaffold. Most LLPS-centric databases analyzed here do not distinguish between drivers and co-drivers, therefore in the rest of the article, when we refer to drivers, we actually mean an umbrella term that covers both drivers and co-drivers.

*In vitro*, co-driver activity is apparent if there are more than one interacting macromolecules strictly required for LLPS or when LLPS of one co-driver occurs at a much lower (physiological) concentration in the presence of the other co-driver. *In vivo*, information on the co-localization of the co-drivers in the same puncta is essential (as often RNA is part of RNP particles).

#### Regulator

Often, the LLPS of a driver or co-drivers requires the presence (activity or activation) of an additional protein, which does not physically take part in condensate scaffold formation, though. There are a variety of regulators, such as modifying enzymes, transport proteins regulating cellular localization, or transcription factors promoting the expression of the driver and/or co-driver.

Although not frequently addressed *in vitro*, one may think of a kinase as a regulator catalyzing a critical phosphorylation event required for the phase separation of the target protein. *In vivo*, knocking out or silencing a gene may reveal a regulator, if its product has a profound (positive or negative) effect on LLPS but does not need to be part of the core assembly of driver/co-driver molecules that constitute the scaffold of condensates.

A ***client*** is not required for and does not have an effect on LLPS (unlike a co-driver or regulator), but it can localize to the condensate formed (through direct or indirect interaction with the driver or co-driver).

This behavior can be demonstrated both *in vitro* and *in vivo* by co-localization of the client with the drivers/scaffolds inducing LLPS. In *in vitro* partitioning experiments, the ability of proteins to enter the condensates already (pre-)formed by other proteins are tested. Importantly, the ability to enter pre-formed condensates can confirm the role as a client, but not as a driver/co-driver.

### 2.2 Four LLPS databases

In the following we briefly outline the main characteristics of the four wider-scope LLPS databases (DBs), whose data are used to derive a consolidated dataset of human proteins that act as drivers/scaffolds in liquid-liquid phase separation processes.

PhaSepDB is a comprehensive resource storing the proteins only on the basis of localization to MLOs [40]. The DB stores information on a total of 2957 proteins. It accepts three different evidence types for protein localization: literature evidence, UniProt localization annotations and the results of high-throughput protein localization experiments. Consequently, PhaSepDB does not categorize the proteins by their role in LLPS.

LLPSDB stores almost 1200 *in vitro* LLPS experiments as entries, addressing the LLPS behavior of 273 proteins [38]. These comprise natural proteins as well as artificially designed protein constructs. The DB provides detailed information on the molecular components and measurement parameters for each experiment, along with their outcomes (e.g. LLPS is detected or not). There is no attempt at interpreting experimental data and/or defining the role of proteins in LLPS.

DrLLPS classifies LLPS-related proteins into scaffolds, regulators and clients as assessed from related literature quotes processed by automated text mining, followed by curator assessment [39]. The underlying literature evidence mostly reports on high-throughput and low-throughput experiments addressing the physical or functional association of proteins with MLOs (for clients), phenotypic effects of their knockout, silencing or overexpression on MLOs (for regulators) as well as on dedicated LLPS experiments (for scaffolds). The DB also includes proteins in different organisms predicted to phase-separate by homology transfer (based on sequence homology with proteins experimentally shown to do so).

PhaSePro stores a relatively small, manually curated set of 121 LLPS proteins as entries, all categorized as ‘drivers’ based on experimental evidence from *in vivo* and/or *in vitro* studies [37]. Rigorous curation criteria are used to categorize a protein as an LLPS driver, which take into account the physiological relevance of conditions reported for the associated experiments. PhaSePro entries also contain information on additional determinants and regulators of the LLPS process as well as on the associated molecular mechanisms.

While two of the four investigated DBs rely on localization/association evidence (PhaSepDB completely, DrLLPS partly), the other two solely rely on dedicated LLPS experiments (LLPSDB and PhaSePro). With regards to the assignment of the specific roles of proteins in the LLPS process, DrLLPS is the only database classifying proteins into distinct categories according to their role in phase separation (scaffolds/regulators/clients). PhaSePro evaluates these roles in the curation process, but only annotates *drivers* as entries. In LLPSDB, which is entirely dedicated to *in vitro* experiments, the role of the protein in LLPS is not explicitly assigned but may be derived from the annotations associated with individual experiments. PhaSepDB contains no information on the role of the protein in phase separation. Although these differences make the integration of the underlying LLPS data rather difficult, here we make an attempt to assess the reliability and level of agreement between the four LLPS resources and integrate their contents.

### 2.3 Data consolidation across LLPS databases reveals inconsistencies in protein annotations

A useful means for assessing the quality and consistency of the data stored in different databases covering a common field of research is to compare equivalent data items across the DBs, evaluate the level of consensus (overlap) and complementarity (differences) in these items, and identify the origins of detected discrepancies [43,44]. We followed this strategy for the four LLPS-dedicated databases. To be able to compare equivalent subsets of the proteins, we restricted our comparison to LLPS drivers/scaffolds, which are considered as synonymous and involve both self-sufficient drivers and co-drivers.

We obtained all the entries, 121 driver proteins from PhaSePro. From DrLLPS only entries labelled as ‘reviewed scaffolds’ have been accepted as drivers, resulting in 150 proteins. Extracting this information reliably from LLPSDB and PhaSepDB, which do not explicitly annotate protein roles, was less straightforward. Using a specific definition and some filtering steps (see ***Methods section 3.1***, and **Figure 1** for details), we extracted 153 native proteins from LLSPDB that could be accepted as LLPS drivers according to our definition, at least regarding their partner dependencies. For the lack of any annotation on their actual role in LLPS, from PhaSepDB we extracted 689 proteins annotated to localize to MLOs by any of the two evidence types of higher confidence, i.e. literature evidence and UniProt localization annotation (**Figure 1**).

**Figure 1:**
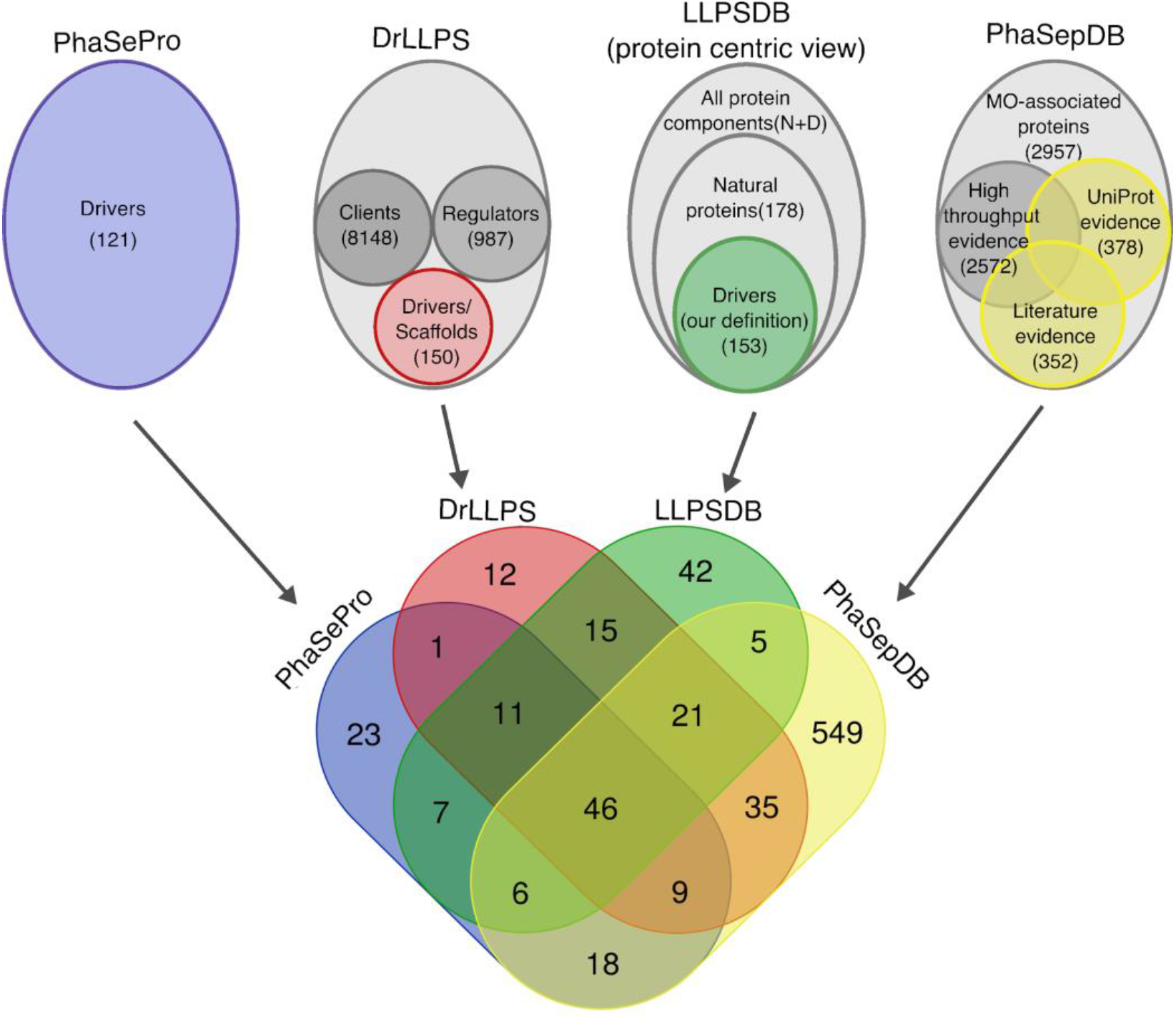
Content and overlap of the four LLPS datasets. The contents of the four databases are represented by ellipsoid shapes in the upper part, with subsets of the data used in this analysis highlighted in color, whereas the rest of their data are depicted in different shades of gray. The total number of proteins as well as the number of proteins in the different subsets are indicated. In the Venn diagram at the bottom, the overlap of the colored subsets of the four LLPS databases is shown.

The overlap of the collected driver proteins across the 4 DBs is in general quite poor (**Figure 1**). In total, only 46 driver proteins are shared by all 4 DBs. PhaSePro, DrLLPS and LLPSDB, taken pairwise, share on average ~54% of their driver proteins. Although the 689 potential driver proteins extracted from PhaSepDB cover ~63% of the other databases’ drivers on average, they are only poorly covered by the remaining 3 DBs taken together (140 proteins; ~20%). This means that although the set of proteins derived from PhaSepDB has a good overall coverage of drivers, it probably also contains many regulators and clients. This lack of specificity apparently stems from the localization-centric rather than LLPS-centric view of PhaSepDB (also resulting in a much larger literature compilation (**Supplementary Figure S1A**) than those of the other resources). Due to the resulting lack of information on the precise role of entries in LLPS, the extraction of driver proteins from PhaSepDB is not feasible, we therefore decided to limit our analysis to the remaining three databases.

**Figure 2A** and **Supplementary Table S1** depicts the overlap of the driver proteins collected from these, PhaSePro, DrLLPS and LLPSDB. A total of 57 driver proteins are shared by all 3 DBs, representing <50% of the proteins annotated as drivers in any of them. A larger number of such proteins are shared by pairs of databases (93 for DrLLPS/LLPSDB, 70 for PhaSePro/LLPSDB, and 67 for PhaSePro/DrLLPS). On the other hand, as many as 135 driver proteins are unique to one of the three DBs with 41, 47 and 47 such proteins found in PhaSePro, DrLLPS and LLPSDB, respectively.

**Figure 2:**
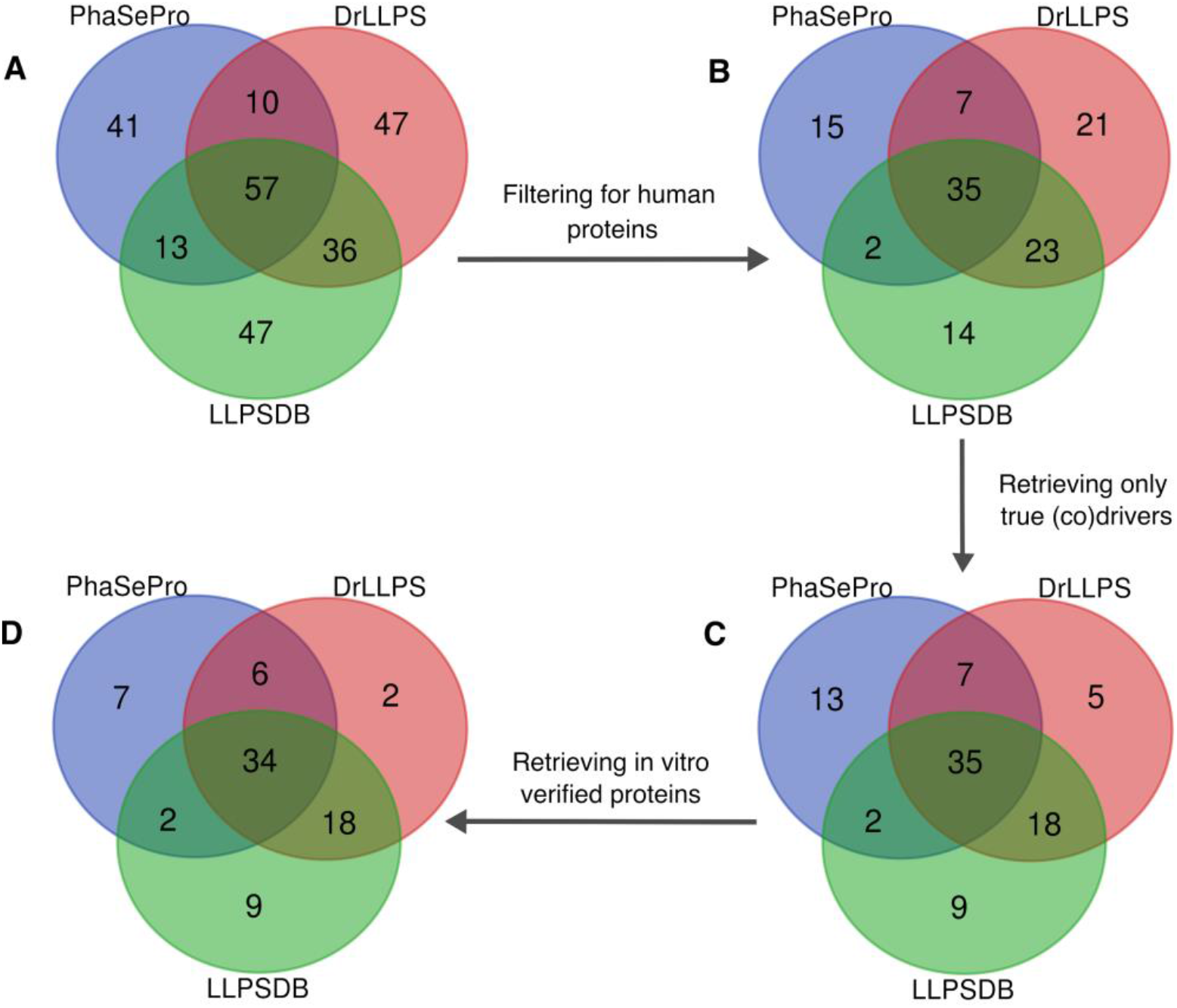
Filtering steps for obtaining a consolidated set of true human LLPS drivers. (A) Venn diagram representing the overlap of the three LLPS databases after the exclusion of data in PhaSepDB. (B) Venn diagram representing the same overlap after filtering for human proteins (117 proteins). (C) The integrated set of 117 human proteins was further filtered for true LLPS (co)drivers by manual re-curation resulting in a set of 89 proteins. (D) Among these we identified 78 in vitro verified proteins, which could be used in the concentration analysis.

Such discrepancies between annotations in databases are not uncommon and may have a number of origins. They may result from (1) differences in the literature that is being covered, (2) differences in curation policies (extracting information from abstracts or full publications, with or without consulting figures and supplementary information), or from (3) making different choices about the information to be archived. (4) Differences in the interpretation of published information may be another important source of observed discrepancies.

With regards to literature covered by the different DBs, each of them curated a unique set of publications, not covered by the other two DBs (**Supplementary Figure S1B**). The reasons for this discrepancy are diverse. LLPSDB stores one reference only, the primary research article, for each of its annotated *in vitro* LLPS measurements. Although the online interface of DrLLPS references several articles for most annotated proteins (both LLPS and localization evidence), the downloadable database file only contains a restricted subset of references enlisted in the “minimum set of experiments” section of the entry pages. The entry pages of PhaSePro also often collect multiple references (including reviews) that describe LLPS or LLPS-related structure-function relations. To gain a similarly restricted set as for the other resources, here we only considered the primary reference for the annotated LLPS region for each entry. The number of unique publications covered by PhaSePro, DrLLPS and LLPSDB takes up 17.8%, 12.4% and 28.7% of the total number of publications covered by all three databases (202), respectively. The fraction of unique publications is the highest for LLPSDB due its experiment-centric rather than protein-centric approach, which ensures that highly-studied proteins (e.g. FUS) are represented by multiple articles. A closer scrutiny also reveals that most of the 41 LLPS drivers unique to PhaSePro, such as TIAR-2, MORC3, U2AF65, Matrin-3 and others are derived from articles published during the summer of 2019, too recent to be considered by the analyzed first releases of the other two databases.

Another source of the detected differences concerns alternative co-driver proteins that can act as substitutes of each other in the same LLPS system. This is for instance the case for, the proteins Gads (the GRAP2 gene product) and SLP-76 (LCP2 gene product), identified as alternatives to Grb2 and Sos1 respectively, in the LAT signalosome [45], the many R-motif containing proteins identified as possible alternative co-drivers acting together with nucleophosmin (NPM1) [46], or the androgen receptor (AR), identified as an alternative of DAXX in driving phase separation with SPOP [47]. These alternative co-drivers are mentioned in the entry pages of the respective systems in PhaSePro, but are not included as independent entries in the database, a direct consequence of the current schema design of this database.

Different curation policies provide another source of discrepancy. For instance, PhaSePro only focuses on native proteins as drivers in experiments carried out under physiological conditions, with DrLLPS and LLPSDB being less strict in this regard. For example, the E3 sumo-protein ligase PIAS2 and tyrosine kinase ABL1 are annotated by DrLLPS and LLPSDB, although the respective experiments were carried out with artificial protein constructs containing 10 tandem copies of the SUMO-interacting motif (SIM) of PIAS2, or 4 tandem copies of the proline-rich region of ABL1 [48]. These constructs introduced artificial, non-physiological multivalency into the studied systems. In accord, PIAS2 and ABL1 are not annotated by PhaSePro, because the native protein chains were not shown to undergo LLPS [48].

Other examples of different curation policies include the yeast RNA-binding protein Mip6, and human γ-D-crystallins. Mip6 is only classified as a scaffold protein in DrLLPS, based on a report that it undergoes LLPS *in vivo* when highly overexpressed [49]. This is only relevant with cellular pathologies associated with concentration imbalance [49], which, however, should not be mixed with annotations of physiological drivers. The annotation of γ-D-crystallins as LLPS drivers by LLPSDB is supported by evidence that rat and human γ-D-crystallins undergo LLPS under high pressure/low temperature conditions, which are only relevant for proteins of deep sea organisms [50]. One can argue that these reports are not representative under physiological conditions, i.e. these proteins should not be accepted as LLPS drivers (see driver definition in ***Results section 2.1***).

An even more fundamental reason for the limited overlap of LLPS drivers in the different databases stems from inconsistencies in interpreting experimental data by database curators. For example, DrLLPS curators appear to employ more lenient criteria for scaffolds compared to those by PhaSePro for drivers, which, in principle, are equivalent categories. For instance, among the entries unique to DrLLPS, we found several proteins (CIRBP, CPEB2, RBM3 and others) that are not demonstrated to undergo LLPS on their own or with a well-defined set of co-drivers. These proteins partition into condensates formed by other proteins [51,52], which defines them as clients but not drivers (cf. ***Results 2.1***).

In other instances, DrLLPS annotates proteins as scaffolds (such as G3BP1, RBFOX1, LSM4, pgl-1 and others), but have borderline evidence only supporting their driver roles by PhaSePro, and thus were annotated as candidate entries. While the central role of G3BP1 and pgl-1, in the formation of the respective liquid-like MLOs (stress granules and *C. elegans* P-granules, respectively) is irrefutable, these proteins were not shown to undergo LLPS *in vitro*, and therefore their partner dependencies are not (yet) sufficiently elucidated.

Another reason for doubting the driver role of certain proteins may lie with the physical properties of the resulting assemblies. For example, human RBFOX1 [53] and yeast LSM4 [54] were not shown to form liquid-like droplets, but were found to assemble into fibrous aggregates of irregular shapes in *in vitro* experiments, and were therefore not classified as LLPS drivers in PhaSePro (see ***section 2.1*** on evidence supporting the liquid material state of condensates). Some of these divergent approaches to the interpretation of the experimental data stem from the complexity of the analyzed systems, and hinge more on curation policies related to this complexity.

### 2.4 Arriving at a consolidated dataset of true human LLPS drivers

Altogether 251 suggested driver proteins could be extracted from the three resources (**Figure 2A**). When filtering for human proteins (this filtering step was required because the following analyses could only be performed on human proteins), 117 drivers remained (**Figure 2B**). We then excluded 28 human proteins that could not be accepted as LLPS (co)drivers/scaffolds according to our definition of this role (see ***section 2.1***), many of which were mentioned in the previous section (**Figure 2C**). In brief, we aimed at retaining only human LLPS drivers where a physiologically relevant form of the protein was studied under physiologically relevant conditions in experiments that have the potential to justify LLPS (co)driver roles (partitioning experiments not accepted), and where the liquid state of the resulting condensates were also confirmed. The comprehensive list of gene names of the excluded proteins (together with reasons of exclusion) is detailed in ***Methods section 3.2***. In our view, the resulting filtered set of 89 human LLPS driver proteins consolidated from the 3 DBs (**Figure 2C; Supplementary Table S2**) is a valuable result on its own that future studies analyzing different aspects of LLPS could also benefit from. Also, it provides an excellent opportunity to further interrogate the underlying data.

### 2.5 Concentrations of LLPS driver proteins

Protein concentration is a crucial parameter of condensate formation manifested in phase diagrams in most LLPS studies. While protein concentrations are only imprecisely quantified or set when probing LLPS inside cells, they can be precisely controlled in *in vitro* LLPS studies. However, an important issue with their interpretation is that we can hardly judge how well they reflect cellular protein concentrations.

To find out if and how this challenge is handled in the field, we compared the concentrations of our confident set of human LLPS drivers used in *in vitro* LLPS experiments to their cellular abundances measured in mass spectrometry (MS)-based quantitative proteomics studies, obtained from PaxDb [55]. To this end, we first filtered our dataset for human proteins with available *in vitro* LLPS measurements and concentration values (**Figure 2D**). For the resulting 78 proteins, the concentrations applied were obtained from LLPSDB and from the literature. We preferred concentrations of full-length proteins over those of segments, and only experiments on native protein forms or those with physiologically relevant modifications (PTMs) or mutations (e.g. phosphomimetic) were considered. Whenever available, multiple concentration values were obtained, and considerable effort was made to obtain the value of saturation concentration (Csat), i.e. the protein concentration minimally required for LLPS to occur, from phase diagrams in LLPSDB [38] or in the respective original publications (see the assembled dataset in **Supplementary Table S3**). PaxDb tissue-specific and cell-line integrated abundance values were also retrieved, converted to concentrations using a published formula [56] and plotted for the same set of proteins for comparison (for technical details on obtaining and converting abundance values, see ***Methods section 3.3***). All obtained concentration values grouped by sources and proteins are provided in **Supplementary Table S4**.

The results of this comparison, depicted in **Figure 3**, show that protein concentrations used in *in vitro* LLPS experiments are in many cases considerably higher, often by one or two orders of magnitude, than cellular concentrations in different tissues or the respective value integrated across different human cell lines. Extreme experimental concentration values were found, for example, in the case of RBM14 [57], NUP153 [58], CCNT1 and DYRK1A [59], where the authors used lyophilized protein powders and provided the applied amounts as mg/ml. For three proteins (LAT, SYN2 and YTHDF3) no PaxDb concentration data could be obtained, hence this comparison could not be made. Averaging all applied concentrations makes no sense since only the lowest concentration presumed to correspond to Csat can be used to judge if the given protein could undergo LLPS under physiologically relevant conditions. Averaging cellular concentrations is also senseless since many of these proteins only detectably express in one or a few specific tissues or cell types, i.e. their abundance in other tissues may be negligible and irrelevant. Due to these reasons, here we compared the lowest *in vitro* concentration where a given protein phase separates (ideally, Csat) to the highest of its PaxDb-derived cellular concentrations (see **Supplementary Figure S2**).

**Figure 3:**
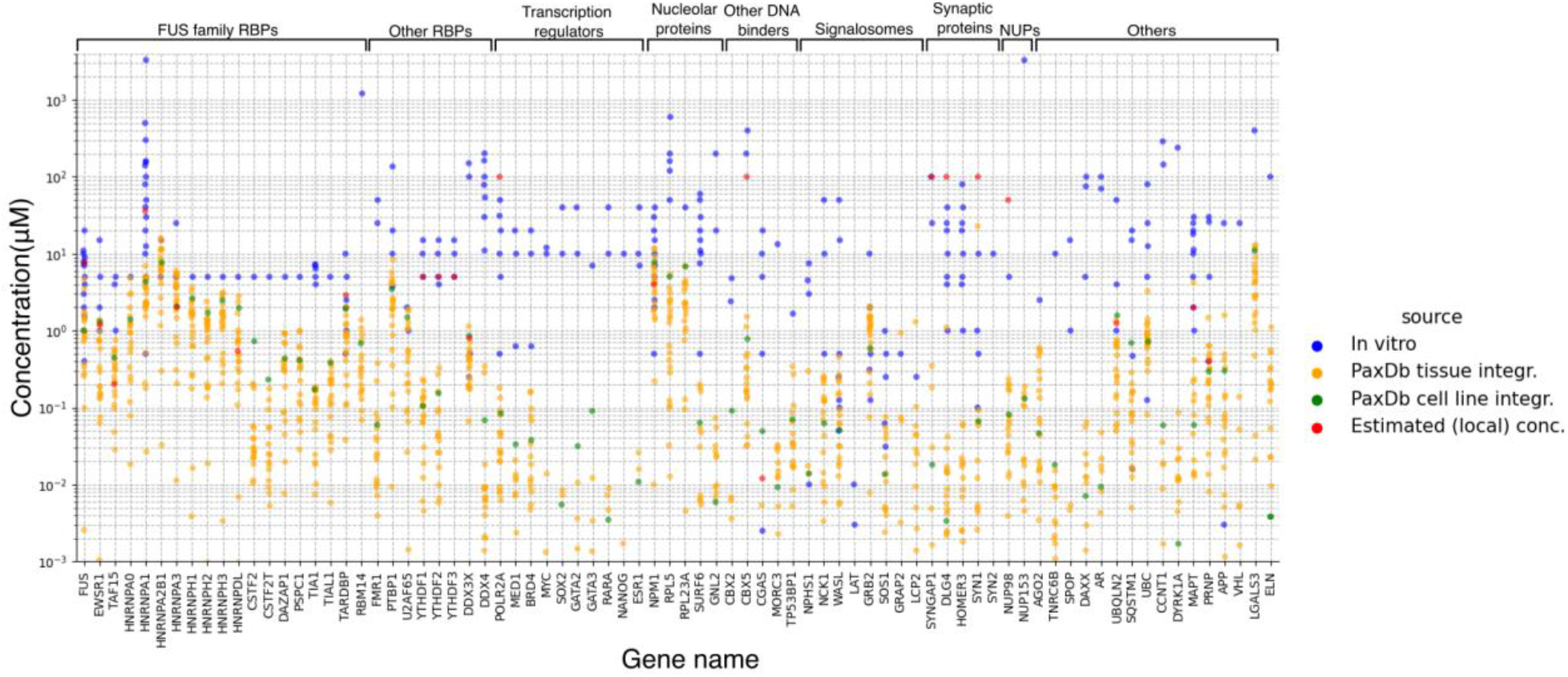
Protein concentrations applied in in vitro LLPS experiments frequently exceed those calculated from proteomics-derived cellular protein abundances. Comparison of protein concentrations applied in in vitro LLPS experiments (in blue), concentrations calculated from proteomics-derived cellular protein abundances stored in PaxDb (tissue-specific integrated values in orange and cell-line integrated values in green) and local concentrations estimated or referred to by authors (in red) for human LLPS driver proteins integrated from the three LLPS resources and filtered for in vitro verified true LLPS (co)drivers. Concentrations are provided in micromolar units, while LLPS proteins are represented by the respective gene names on the horizontal axis. Functional groups of the proteins are indicated on the top of the figure.

By this comparison, only 25 of the 75 proteins (33.3%) could undergo LLPS *in vitro* at a concentration that is within the range of its PaxDb-derived cellular concentrations. A further 17 proteins (22.7%) could undergo LLPS *in vitro* at a concentration that is within one order of magnitude from its highest PaxDb-derived cellular concentration. The remaining 33 proteins (44%) either could not undergo LLPS at close-to-cellular concentrations or they haven’t even been tested. It is important to mention here that judging these relations is also difficult because *in vitro* experiments were often carried out only at a single “comfortable” concentration. That is, no concentration-dependent phase diagram was reported for at least the following 30 proteins (represented by gene names), therefore their saturation concentrations were not defined and thus their ability to undergo LLPS at physiologically relevant concentrations can hardly be judged: NUP153, RBM14, LGALS3, CCNT1, DYRK1A, CSTF2, CSTF2T, HNRNPDL, HNRNPA0, HNRNPA2B1, HNRNPA3, HNRNPH1, HNRNPH2, HNRNPH3, TIAL1, DAZAP1, PSPC1, SOX2, RARA, NANOG, MYC, GATA2, ESR1 VHL, ELN, RPL23A, NUP98, GATA3, LCP2, SYN2. This partly explains the concentration deviations observed, since these proteins take up more than half of the groups where the applied concentrations show no overlap with the range of cellular concentrations (9/17 - 53% and 18/33 - 54.5%, respectively). Further, for some of the investigated proteins the authors calculated or referred to previously measured/estimated local concentrations (marked as red dots on **Figure 3**) that are often orders of magnitude higher than the respective PaxDb-derived concentrations, to justify the concentrations applied in their experiments (see **Table S4** for these data).

As these large systematic differences are alarming with regards to the dependability of *in vitro* LLPS experiments and the phase-separating potency of the proteins studied, we next examine their potential origins and discuss the possible implications with regards to *in vitro* LLPS studies, current estimates of protein cellular concentrations, and the organization of the cellular proteome.

### 2.6 Cellular protein concentration: an elusive parameter

Since LLPS is highly concentration-dependent, the finding that *in vitro* concentrations applied in LLPS experiments are apparently higher than the ones estimated for cells seems perplexing at first glance. However, there are several considerations that help resolve the associated concerns, and thus provide grounds for optimism regarding most LLPS experiments.

#### 2.6.1 The uncertainties associated with current cellular protein abundance measures

First, we note that cellular protein concentrations analyzed here are derived from abundance determined in quantitative proteomics studies. These values in PaxDb represent the relative number of protein copies expressed in units of parts per million, and are converted to protein concentrations by published estimates of the number of protein molecules/fl cellular volume in model organisms, including human, and the average estimated cell volume [56] (see ***Methods*** for detail).

One weakness of proteomics methods is that they are biased against lowly expressed proteins and membrane proteins, which tend to be underestimated [60–63]. This is especially true for membrane proteins that also suffer from solubility issues [64]. Solubility issues may also hamper accurate detection/quantification of proteins that are part of supramolecular complexes or condensates [65], however this possibility still needs to be explored. By comparing the cellular abundance of LLPS drivers to the whole human proteome, LLPS drivers tend to be of low abundance (**Figure 4**). This is not surprising as many of the proteins driving LLPS belong to protein families typically of low abundance, such as signaling proteins [62], transcription factors [63,66], chromatin-associated proteins [67] or postsynaptic density proteins [68,69], being at the detection limits of proteomics studies. We may thus reasonably assume that cellular protein concentrations converted from PaxDb abundance data in **Figure 3** are, in general, underestimated for LLPS drivers.

**Figure 4:**
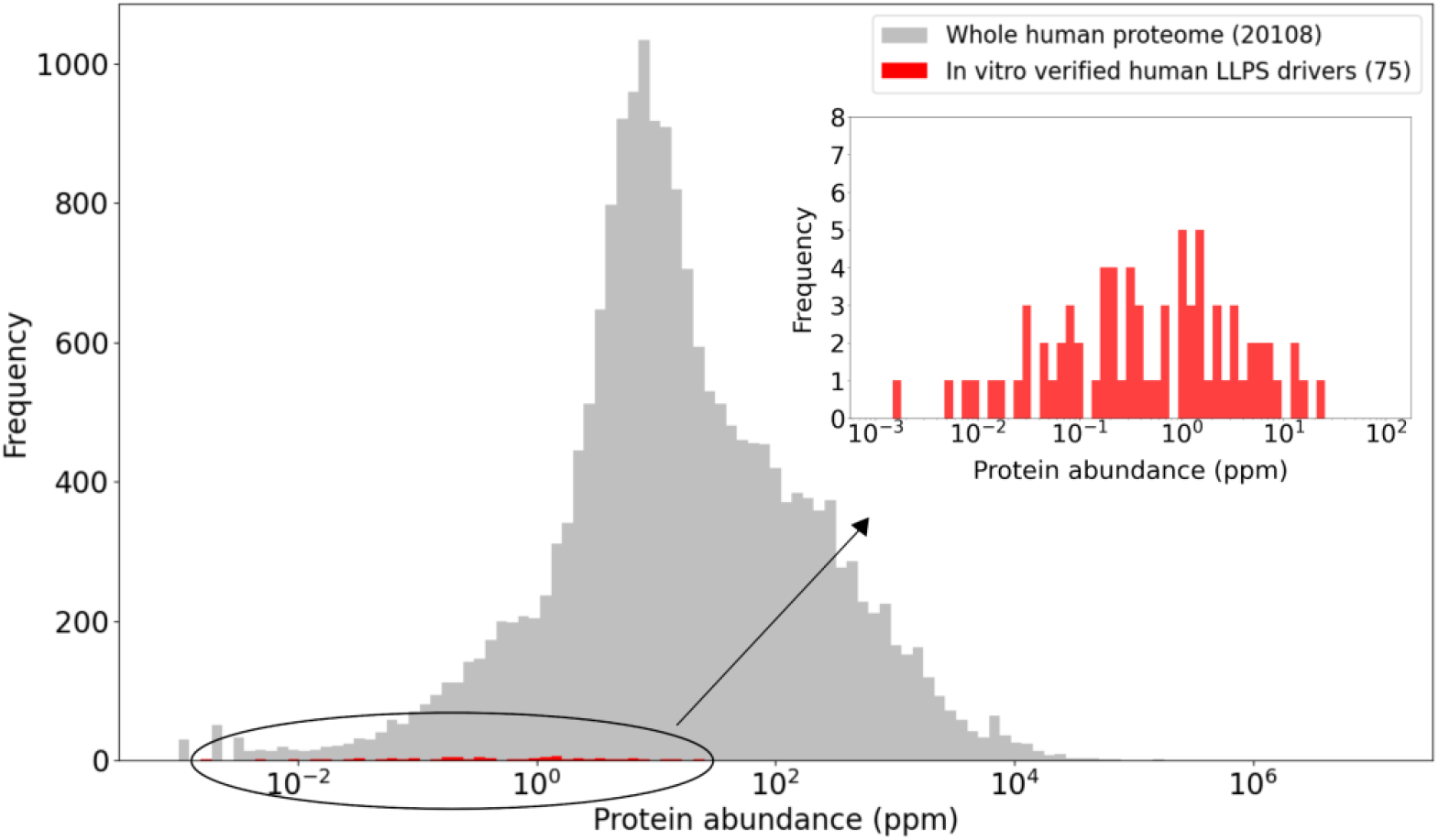
Human LLPS driver proteins are of relatively low abundance compared to the proteome average. The highest tissue- or cell type-specific integrated abundance value was derived for each human protein from PaxDb. These maximal abundances of human LLPS driver proteins (red) are compared to those of the proteome (grey) using histograms. The abundances of LLPS drivers are also separately depicted in the inset histogram to ensure better resolution of the data. Abundances are provided in parts per million (ppm) units on the X axis.

#### 2.6.2 Local concentrations of proteins in their cellular niche define their phase diagrams, but what about measuring them?

Another potential error leading to the underestimation of actual protein concentrations may arise from the conversion of protein abundances to cellular concentration values. On the one hand, the applied formula [56] uses an average cell volume that can lead to a significant error if the protein is mostly expressed in cell types much smaller or larger than the average. On the other hand, the formula assumes that proteins are evenly distributed in the entire volume of cells, which is probably never the case. Most proteins are not evenly dispersed within cells but are localized to specific cellular niches, which define the specific biochemical environment and sets of interaction partners, required to carry out their function [70]. While a number of mass spectrometry-based techniques, referred to as *spatial proteomics*, are able to reliably assign the subcellular localizations of thousands of proteins and confirm the above tendency, their ability to quantify protein copy numbers in specific localizations is still limited [71,72]. For the lack of other dedicated techniques, we need to accept that *in vivo* local protein concentrations are currently impossible to precisely quantify. However, it is important to realize that the discovery of LLPS and MLOs brought us closer to understanding how cellular material is organized. Such specialized liquid condensates (as well as classical membrane-bound organelles) enrich some, but exclude other, proteins. The more organelles or condensates are discovered, the less accurately proteomics-derived cellular concentration measures represent actual, effective protein concentrations due to “diluting” proteins to whole cell volumes.

#### 2.6.3 Molecular mechanisms facilitating asymmetric/spatially restricted protein distributions

There are at least two molecular mechanisms that ensure high local concentrations of even low-abundance proteins. Although many proteins are transported to their respective cellular niches after translation, asymmetric protein distribution can arise by local translation after mRNA localization. This is usually mentioned in conjunction with neurons, in the context of the transport of mRNAs to axon termini followed by the local translation of proteins functioning in synaptic transmission [73,74]. However, it was also described for migrating fibroblasts [75] and it is reasonable to assume that local translation is a general mechanism in most cell types. It is also important to emphasize here that proteins rich in intrinsically disordered regions with the potential to promote the formation of reversible cellular assemblies show an increased propensity for local translation, most probably to ensure interaction fidelity and signaling sensitivity [76]. Among our studied LLPS driver proteins, local translation reportedly applies to postsynaptic-density proteins in nerve terminals, such as PSD-95 (gene DLG4), SynGAP (gene SYNGAP1), Homer-3 (HOMER3) and Synapsin-1 (gene SYN1), which reach local concentrations of ~100 μM in quantitative proteomics studies of purified postsynaptic densities (PSDs) and nerve terminals [65,77,78]. These high local concentrations entirely justify respective *in vitro* LLPS measurements [79,80] and invalidate PaxDb-derived negligible cellular concentrations (1.1 μM, 0.35 μM and 0.062 μM for DLG4, SYNGAP1 and HOMER3, respectively).

Another potentially general process that may ensure high local concentrations is high-affinity anchoring to sites/surfaces that display specific binding sites in large numbers. In such cases, local concentrations of the proteins can be defined by the occupancy and spatial arrangement of binding sites, the former being influenced by multiple factors, including concentration in the bulk phase, binding affinity, k_on_ and k_off_ rates, etc. Multidomain, modular LLPS drivers are often specifically anchored to RNA, DNA or membranes that display multiple repeated binding sites for one of their domains, while their phase separation is usually mediated by disordered regions of low sequence complexity located outside those domains. To cite a few known examples among our LLPS drivers, the U2AF65 splicing factor is anchored to long pyrimidine-rich RNA regions via its RNA-recognition motifs (RRMs) [81]; YTHDF proteins are anchored to polymethylated mRNAs through their YTH domains [82]; VHL is anchored to long dinucleotide (CU or AG) repeats of stress-induced noncoding RNAs via a short cationic segment [83]; Pol II is anchored to the transcribed DNA segment and the transcription machinery in high numbers, reaching high local concentrations [84]; HP1α is anchored to DNA and specific histone marks as well as MORC3 [85]; and ERα is anchored to DNA with closely adjacent estrogen response elements via its nuclear receptor DNA-binding domain [86]. Also, FG-rich nucleoporins NUP98 and NUP153 are anchored to the nuclear pore complex (NPC) scaffold through specific domains outside their FG-rich regions, resulting in more than 200x increase in their local concentration [87]. In all, in the above cases the LLPS-prone regions are distinct from the anchoring domains, and high-affinity anchoring is an independent event that ensures that LLPS-prone regions locally accumulate, effectively promoting LLPS. Accordingly, *in vivo* local protein concentrations estimated from binding-site frequencies and occupancies were in some cases used to justify the higher protein concentration range applied in *in vitro* LLPS studies [88].

It was shown, for example, that only the 48 copies of NUP98 that are obligate subunits of the nuclear pore complex (NPC) ensure a local concentration of 50 μM (2.5 mg/ml) of the FG domain and 2 mM of the constituent FG motifs. When considering other FG NUPs as well, this lower bound increases to 6 mM for FG motifs [87]. The fact that the high local concentrations derived from anchoring (from binding site occupancies and frequencies) are mostly closer to the protein concentrations measured inside condensates than those in the bulk phase does not mean that they cannot be used to justify high *in vitro* applied concentrations, because anchoring is independent from and upstream to condensate formation by LLPS. This relationship is easiest to understand through the following hypothetical experiment: If NPCs were treated with 1,6-hexanediol to destroy the weak interactions between FG motifs that form the phase-separated dense polymer meshwork within the NPC channel, then the condensates would dissolve and transport selectivity diminish, yet local concentrations of NUPs would still be very high, since 1,6-hexanediol could not destroy high-affinity anchoring of the NUP subunits to the NPC scaffold. Therefore, high local concentrations of LLPS drivers calculated based on binding site frequencies and occupancies should be accepted as a justification for using higher protein concentrations in *in vitro* LLPS studies.

In all, many if not most LLPS-driver proteins are of relatively low abundance but display highly restricted patterns of subcellular localization often due to local translation and/or anchoring, suggesting that their local concentrations can be several orders of magnitude higher than their cellular concentrations derived from proteomics (**Figure 3**). Since the phase behavior of proteins is determined by their effective, local concentrations, *in vitro* experiments should only be expected to respect those to be acknowledged as physiologically relevant.

#### 2.6.4 Other potential reasons for higher concentrations of proteins needed to be applied in LLPS experiments

An important reason behind the use of higher protein concentrations in LLPS experiments could be that they are required to compensate for the lack of right partners and conditions that would promote a more physiological-like LLPS process. This is clearly the case for several transcription factors (namely GATA2, SOX2, MYC, NANOG, RARA, ESR1 [52]) and co-activators (namely MED1 and BRD4 [89]) that had been initially studied separately or just mixed with each-other. The physiological relevance of the results of these preliminary experiments using high concentrations of such low-abundance transcription regulators are strongly questionable. However, a more recent study not yet included in any of the dedicated resources demonstrated that transcription factors and co-activators can indeed undergo phase separation and form enhancer condensates at physiological concentrations in the presence of DNA fragments harboring multiple copies of the specific recognition elements of the transcription factors [90]. In this case, the DNA serves as an anchor point, enabling a molecular arrangement that lowers the saturation concentrations of the proteins. HP1 proteins represent a similar case, they were first studied to undergo LLPS on their own or just with DNA [88,91]. Later, chromatin marks and important protein partners were also discovered that contribute to their LLPS and remarkably lower their saturation concentrations [92].

A further point often missed is that *in vitro* LLPS experiments often just use a representative protein of a larger cohort of similar proteins, and in such cases high applied concentrations of the representative need to compensate the lack of others. This is clearly the case for NUPs, where usually only one NUP, in our case NUP98 [87] or NUP153 [58] is used in the experiments, but several similar FG-rich NUPs actually contribute to the phase-separated dense polymer meshwork formed within NPC channels, exerting a “family effect” *in vivo* [87].

Yet another important factor to consider is the effect of crowding. The presence of even subtle amounts of crowding agents remarkably boosts most LLPS processes hitherto studied *in vitro* [27,93,94]. However, crowding agents used in LLPS experiments (usually 1-12% in volume) fall short of matching cellular crowding conditions, so their boosting effect on LLPS may be far below the effect exerted by the totality of cellular macromolecules [95], which could again justify the need for somewhat higher *in vitro* protein concentrations as a compensation. At the same time, cellular macromolecules could also negatively affect LLPS [96].

In all, there are a number of reasons to believe that cellular concentrations underestimate the effective, local concentrations of proteins in general and even more severely underestimate those of most LLPS drivers in particular. Also, higher protein concentrations may be required to be applied in *in vitro* experiments to compensate for other effects, such as the lack of partners, lack of similar family members or the lack of appropriate crowding conditions. Yet another important evidence supporting the physiological relevance of *in vitro* condensate formation by our selected drivers is that for most of them *in vivo* observations also support that they can form punctate structures/foci/liquid granules within living cells (109 of the 121 PhaSePro proteins have some kind of *in vivo* evidence of LLPS). Actually, *in vivo* observations usually precede and serve as an incentive for downstream *in vitro* experiments. Although it is just a small fraction of the *in vivo* experiments where expression of the labelled protein was driven by its endogenous promoter [97], in most cases a significant overexpression was deliberately avoided even if inducible promoters were used. Furthermore, silencing or truncating the respective driver gene often has led to a complete disappearance or remarkable changes in the phenotypes of the associated MLOs. Due to all the above reasons, we are confident that the fundamental role of our select drivers in the formation/integrity of biologically relevant condensates is mostly well-supported, even if their *in vitro*-applied concentrations tend to exceed proteomics-derived cellular concentration levels. Next, we provide strong evidence for this tenet by observing the dosage sensitivity of genes encoding for these proteins.

### 2.7 Dosage sensitivity of LLPS driver genes

Being a highly cooperative phase transition, the primary determinant of LLPS is protein concentration [46,98–100]. If LLPS is an important functional property of our LLPS drivers, their availability in the cell would need to be tightly regulated. It was even suggested that proteins with the ability to trigger LLPS may become toxic upon increased expression, which could even cause dosage sensitivity to lead to disease [49]. Furthermore, LLPS-associated proteins are enriched in IDRs prone to promiscuous interactions, properties that also require tight regulation of cellular concentrations of proteins [101] and dosage sensitivity of the corresponding genes [102]. Gene dosage is defined as the copy number of a particular gene in a genome, and dosage sensitivity is the measure of intolerance to modifications of gene dosage. So far the possible relationship of LLPS-associated proteins with dosage sensitivity was limited to a comparison of predicted physicochemical properties (disorder, RNA-binding, amino acid composition) of proteins that become toxic upon overexpression in yeast with those that localize to granules [49].

Therefore, here we use our consolidated dataset of human LLPS drivers to clarify the possible relationship of LLPS and dosage sensitivity. To this end, we evaluate the extent to which our 89 human LLPS drivers (**Figure 2C**) are over- or under-represented in the sets of the most reliable dosage sensitive (MRDS) and most reliable dosage insensitive (MRDIS) human genes recently consolidated [103].

We found a strong enrichment of human LLPS-associated genes in MRDS genes (chi^2^ test, p<0.00001) (**Figure 5A**) and a strong depletion in MRDIS genes (chi^2^ test, p=0.000221) (**Figure 5B**) using the reviewed human proteome from UniProt [104] as a background. We noted that human LLPS proteins are generally very well annotated, possibly much better than non-LLPS proteins. To avoid bias from the different levels of annotation among human proteins, we repeated the analysis by using randomized selections of similarly well-annotated subsets of the human proteome as a background. LLPS-associated genes displayed a much higher overlap with MRDS genes and a much smaller overlap with MRDIS genes than any of the equivalent 1000 random gene sets, providing solid statistical evidence for their dosage sensitivity (**Figure 5AB**).

**Figure 5:**
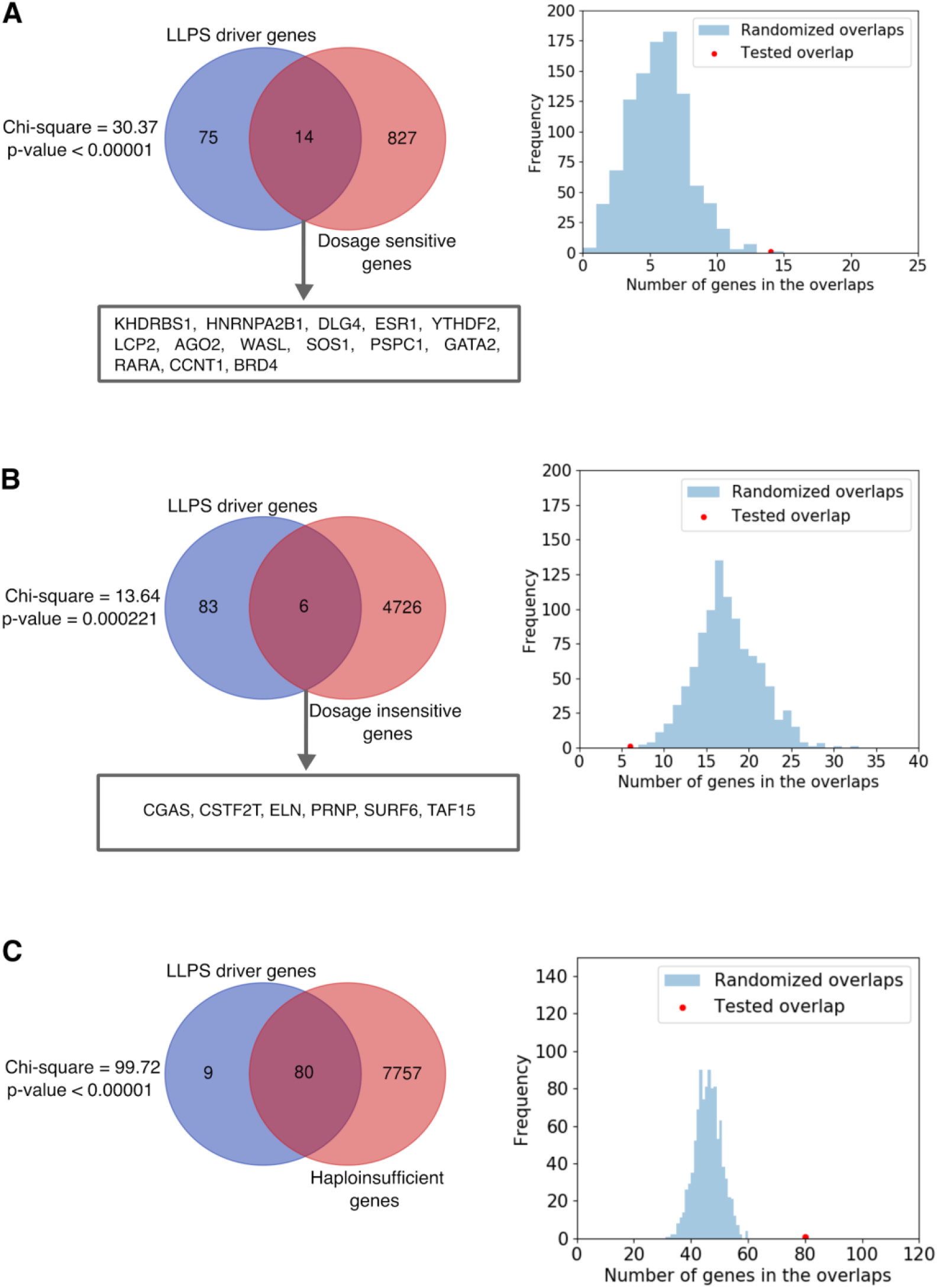
The genes of human LLPS driver proteins are overrepresented in dosage sensitive genes. The Venn diagrams (left) show the overlap between human LLPS driver genes and the set of most reliable dosage sensitive (MRDS) genes (A), the set of most reliable dosage insensitive (MRDIS) genes (B) and the set of haploinsufficient genes (C). The genes in the intersections are listed for A and B. The detected overlaps represent statistically highly significant enrichments (A and C) or depletion (B) based on chi^2^ tests using the whole human reviewed UniProt proteome as background. Histograms (right) showing the difference between the overlaps of human LLPS-associated genes (red) or equivalent sets of randomly selected well-annotated genes (blue) with (A) MRDS, (B) MRDIS and (C) haploinsufficient genes.

The set of dosage-sensitive genes [103] was consolidated from 4 main sources, comprising 2 datasets of haploinsufficient genes [105,106], and one dataset each of ohnologs [107] and copy number-conserved genes [108], which display different flavors of dosage sensitivity. Haploinsufficiency measures the intolerance to heterozygous loss of function (when protein products of one of the two alleles are lost) [103,105]. Ohnologs are pairs of genes originating from whole-genome duplications [107]. If a gene has only ohnologs but no paralogs, or shows conserved copy numbers across mammalian genomes, it is also considered as dosage sensitive [103]. To find out which of these properties contribute to the detected dosage sensitivity of LLPS-associated genes more, we retrieved the above-mentioned 4 original datasets, mapped them to UniProt, integrated the two datasets of haploinsufficient genes and performed the relevant enrichment analyses (for the integrated datasets, see **Supplementary Table S5**). We found that LLPS-associated genes are highly enriched in haploinsufficient genes (80/89 were found in the list of 7837 haploinsufficient genes) (**Figure 5C, Supplementary Table S6**). LLPS-associated genes were also enriched in ohnologs (**Supplementary Figure S3A**) but not in copy number-conserved genes (**Supplementary Figure S3B**) based on chi^2^ tests. These results were confirmed by analyses based on randomized selections (**Figure 5C and Supplementary Figure S3**, **Supplementary Tables S6 and S7**).

The MRDS dataset [103] is significantly enriched in transcription factors (TFs), which tend to be tightly regulated low-abundance proteins. To verify that the observed highly significant enrichment for haploinsufficient genes or ohnologs is not due to an enrichment of LLPS drivers for transcription factors, we evaluated the enrichment for TFs in the LLPS-associated dataset, using the same dataset of transcription factors as in [103] (a list of 1639 TFs originally published in [109]). The results showed no enrichment in TFs among the LLPS-associated genes (9 TFs of 89 genes; chi^2^ test, *p*=0.456 using the reviewed human proteome as background), as well as no enrichments in TF’s among the LLPS-associated genes that overlap with the MRDS, haploinsufficiency or ohnologs datasets as shown by **Supplementary Figure S4.**

This analysis indicates that both losses and gains in the copy number of LLPS-associated genes may be deleterious to cells, i.e. their dosage sensitivity is not an exclusive consequence of the overexpression of the corresponding gene products as previous suggested [49], but reflects a deleterious perturbation of the cellular availability of LLPS-associated proteins, in line with their important roles. In our view, the propensity for dosage sensitivity reflecting a tight regulation of their availability lends strong support for their role in *in vivo* condensate formation, validating the results of associated *in vitro* LLPS experiments.

## 3. Methods

### 3.1 Analysis of the overlap between LLPS databases

In order to analyze the overlap between the 4 publically available LLPS databases, we needed to make sure that we compare the subsets of their data that are equivalent to each other. PhaSepDB is based on protein localization evidence (association to MLOs) and not on the ability to undergo LLPS, so it was not possible to filter for LLPS drivers. Therefore, only the more confident UniProt annotated and reviewed subsets of the database were obtained from PhaSepDB [40] (version 1.3, October 2019), entries based on high-throughput evidence were not included. From PhaSePro [37] (version 1.1.0) we obtained all the 121 entries. DrLLPS [39] (version 1.0) is represented by its 150 reviewed scaffold proteins in this analysis. From LLPSDB [38] we only accepted natural proteins that have shown phase separation in at least one of the corresponding experiments. Since this database stores *in vitro* LLPS experiments as entries and does not categorize the investigated proteins according to their role played in LLPS, we needed to introduce a definition of drivers based on the presented *in vitro* experiments. We accepted those proteins as drivers that could undergo phase separation in any of the associated experiments on their own, as one-component systems. From the other subset of proteins that could only undergo LLPS together with other main components (two- or multi-component systems with other proteins, DNA or RNA), we only accepted those as (co)drivers that were essential for LLPS to happen in the given experiment, meaning that the rest of the main components involved in the particular experiment could not undergo LLPS on their own in another experiment. This filtering was necessary in order to make sure that proteins used as accessory components (like EGFP or mCherry) or regulators in the experiments are not accepted as drivers. We obtained 153 proteins from this filtering. However, we note that our simple definition applied for automated filtering of the data only considers partner dependencies and could obviously not sufficiently substitute for the considerate evaluation and interpretation of experiments by database curators (for instance, the physiologically relevant nature of experiments could not be judged at this point). Comparison of the entries of the databases (provided as **Supplementary Table S1**) was done based on UniProt ACs of the canonical proteins, information on isoforms (even where available) was omitted.

It is important to mention that for some of the driver proteins the full-length protein has never been tested for LLPS in their respective primary publications, only smaller segments have been expressed and purified. These proteins were also accepted as drivers, based on the assumption that if a segment of a protein is prone to undergo LLPS then most likely the full protein will also be able to undergo LLPS.

### 3.2 Integration and filtering of LLPS driver proteins

Since PhaSepDB does not contain any annotations on what role the included proteins play in LLPS, and the associated literature did not show a good overlap with those of the other three resources, PhaSepDB was excluded from data integration. 251 proteins could be integrated from the other three resources. This set was filtered for human proteins, 117 remaining. In the next step we excluded those human proteins that could not be accepted as LLPS (co)drivers/scaffolds according to our definition of this role.

Proteins under the gene names ABL1, PIAS2, SUMO3, ITSN1 and C9orf72 were excluded because they only served as donors of protein modules amplified into artificial repeat proteins for LLPS experiments and the natural proteins have never been tested for LLPS (see associated experiments in LLPSDB). CRYGD and EN2 were excluded because they were only tested for LLPS under highly non-physiological conditions (high pressure and/or low temperature). TP53, KPNB1, KPNA2, FYN, MAP1LC3B, XPO1, RBM3, CPEB2, CIRBP, SGO1 and FBN1 were excluded because they were only shown to partition into the already formed condensates of other proteins (based on which they could be accepted as clients but not as (co)drivers), but were not demonstrated to undergo LLPS on their own or as necessary components of multicomponent LLPS systems (see associated experiments in LLPSDB and/or DrLLPS). RBFOX1 was excluded because there is no evidence for liquidity of the aggregates it formed [53]. 5HT1A was excluded due to the respective study reporting on concentration units, particularly 1:19 w/v that we could not convert into μM [110]. Although HNRNPAB, HNRNPA1L2 and HNRNPD are present in LLPSDB, they were excluded because we could not see any droplets in the figure panels presenting their respective droplet formation experiments [27]. The above listed 23 proteins were excluded due to insufficient *in vitro* LLPS evidence, while further 5 proteins (namely ELAVL1, G3BP1, DYRK3, AXIN1 and ZNF207) were excluded based on insufficient *in vivo* evidence (proteins supported by only *in vivo* evidence were accepted as drivers if both their localization to liquid foci and their necessity for the formation/integrity of the respective foci were proved). The resulting set of 89 proteins can be found in **Supplementary Table S2**.

### 3.3 Obtaining, converting and comparing protein concentration and cellular abundance values from different sources

We first obtained the protein concentrations applied in *in vitro* LLPS experiments (only those where LLPS was detected) either from LLPSDB (where available) or from the primary publications provided by the other resources. Where available, concentration values for the full-length wild type proteins were used, otherwise concentrations of segments were also accepted. If there were multiple measurements with a positive LLPS outcome (e.g. with different measurement conditions, partners), we accepted all applied protein concentrations where a physiologically relevant form of the protein was studied (including simple truncates, modifications, mutations mimicking modifications) to gain a distribution rather than a single data point. In cases when a concentration range was applied to produce a phase diagram [46,98–100], we first took the average of the extreme values of the range (such averaging was performed for 86 of the 366 data points). In a subsequent manual curation round, these averaged values were replaced by the saturation concentrations if we could find a statement from the authors on the minimal concentration required for LLPS in the respective manuscript, or if it could be read from a concentration-dependent phase diagram provided by LLPSDB or by the original article (see **Supplementary Table S3** where the source is indicated for each datapoint). We accepted all conventional concentration units, like mg/ml, μM, nM and converted them to μM. When converting units in mg/ml to μM, we made sure that we calculate with the molecular mass of the protein construct that was applied in the respective experiment, taking into account truncations of the protein chain, fused proteins (usually GFP), as well as indicated sequence tags. We used the Compute pI/Mw tool of the ExPASy server (https://web.expasy.org/compute_pi/) to calculate the molecular weights of protein segments.

In case of LAT the concentrations were provided as molecules/μm2 as it is a membrane protein, however the authors provided an estimation on what protein concentration this is equivalent to [45,55], so we accepted their estimate and used the same estimate in case of nephrin (gene NPHS1), another membrane protein studied by the same research group.

Proteomics-derived protein abundance values were obtained from the integrated datasets of PaxDb, the Protein Abundance Database (version 4.1) [55]. To this end, UniProt ACs provided by the LLPS databases were mapped onto Ensembl protein IDs and abundance data available for the latter have been retrieved from PaxDb. If there were multiple available (tissue-specific or cell line integrated) abundance values for a protein, we accepted all to gain a distribution of data rather than a single data point. The PaxDb abundance values reported in ppm (parts-per-million) as units were converted to micromolar concentrations using the below formula Eq. (1) from [111], where *k* ≈ 3·10^6^ proteins/fL, the Avogadro constant *N_A_* = 6.02 × 10^23^ molecules/mol, and A is the abundance. This allowed comparisons with the *in vitro* applied LLPS protein concentrations retrieved from LLPSDB and the literature. For some of the proteins, like LAT (transmembrane), SYN2 (membrane anchored) and YTHDF3, PaxDb did not have any abundance values, so we only show the concentrations reported for their *in vitro* LLPS experiments in the concentration graphs without a comparison.

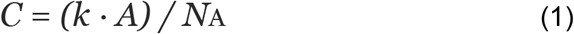

The publications reporting on the in vitro LLPS experiments performed for human LLPS proteins were screened for author statements referring to published or calculated estimates of the physiologically relevant cellular or local concentrations of the investigated proteins. These concentrations were collected (where available) and used as reference values besides PaxDb-derived abundance-based concentrations. For all the obtained concentration values depicted in **Figure 3** grouped by genes and data sources see **Supplementary Table S4**.

For the abundance comparison depicted in **Figure 4** the highest tissue- or cell line-specific integrated PaxDb value was obtained for each human protein with such PaxDb data in ppm (approximately the full proteome) and their frequency distribution was used as reference to be compared to the highest abundances of LLPS driver proteins.

### 3.4 Dosage sensitivity enrichment analyses

The lists of 853 most reliable dosage sensitive (MRDS) and 5579 most reliable dosage insensitive (MRDIS) human genes were downloaded from the supplementary material of Ni *et al.* [103]. The lists of 3230 (probability of being LoF-intolerant (pLI) > 0.9) and 7841 haploinsufficient human genes were obtained from Lek *et al.* [105] and Shihab *et al.* [106], respectively. A list of 7,294 ohnologs was collected from Makino *et al.* [107], while a list of 7014 copy number-conserved genes was taken from Rice *et al.* [108]. The reviewed human proteome (20359 proteins) was retrieved from UniProt release 2020_04 [104]. MRDS, MRDIS, haploinsufficient and copy number-conserved genes, as well as ohnologs were mapped against UniProt by using the provided Ensembl transcript/gene identifiers (where available) or gene names. Only those were retained in the datasets that could be mapped against the reviewed human proteome. After merging the two lists of UniProt ACs for haploinsufficient genes, we gained 841, 4732, 7837, 6865 and 6948 genes for the five properties, respectively. The obtained gene lists with the mapped UniProt ACs are provided as **Supplementary Table S5**. The overlaps of the 89 human LLPS drivers with the five gene sets were computed, and chi^2^ statistics were applied to address statistical significance of the overlaps using the reviewed human proteome (20359 proteins) as background.

Since >95% of our integrated human LLPS driver proteins are reviewed UniProt proteins with an annotation score of 5 of 5 and evidence at protein level, we filtered the reviewed human proteome of UniProt for similarly well-annotated proteins, and used the resulting subset of 13389 proteins as a background for randomized selections to avoid any biases stemming from the large differences in annotation levels of human proteins. 1000 protein sets of equivalent size to the set of 89 human LLPS drivers were selected and their overlaps with MRDS, MRDIS, haploinsufficient and copy number-conserved genes as well as ohnologs were compared to that of LLPS drivers. These data are provided as **Supplementary Tables S6** (for LLPS drivers) **and S7** (for equivalent randomly selected genes sets).

### 3.5 Data analysis and representation software tools

The obtained data was analyzed with custom-made python scripts (Python version 3.7.9). Venn diagrams were produced by the UGent Bioinformatics and Evolutionary Genomics group’s online tool (http://bioinformatics.psb.ugent.be/webtools/Venn/). The histograms and the concentration graphs were generated using Python’s seaborn (version 0.10.1), matplotlib.pyplot (version 3.3.2) and pandas (version 1.0.1) modules.

## 4. Conclusions

Mounting experimental evidence for the phase separation of macromolecules and the formation of MLOs inspired the development of several dedicated data resources. By comparing the data of four wide-scope LLPS resources, we found significant differences in their annotated evidence types, curation policies and interpretation of experimental results [41,42]. By integrating the LLPS driver/scaffold proteins from PhaSePro, LLPSDB and DrLLPS, we generated a comprehensive compendium that we further filtered to obtain a set of 89 human LLPS drivers supported by physiologically relevant experiments.

Since protein concentration is the key factor of LLPS, we used this high-confidence dataset to investigate this parameter in great detail. Our in-depth analysis highlighted a common bias for using higher protein concentrations in *in vitro* experiments than the ones calculated from proteomics data for cells. We show, however, that there is a range of considerations that can resolve these concerns. First, for a large fraction of the analyzed proteins the saturation concentration was not identified and thus only arbitrarily selected concentrations could be used in the comparison. Second, many LLPS drivers are of low abundance and/or membrane-associated, and thus their relative abundance is generally underestimated in proteomics measurements. Third, in the analyzed, often preliminary *in vitro* LLPS experiments higher protein concentrations were sometimes required to compensate for the lack of appropriate partners, the lack of similar drivers (with “family effect”), or the lack of physiological crowding conditions. Fourth, it seems generally difficult – or even impossible – to derive meaningful absolute concentration values from quantitative proteomics studies [71,112], as they can seriously underestimate the effective, local concentration of proteins, especially for those with highly restricted localization like most LLPS drivers. We anticipate that with further improvements in our understanding of cellular organization by membrane-bound and membraneless organelles, the assumption of even protein distributions in the cell becomes increasingly less tenable. The overrepresentation of human LLPS drivers in dosage sensitive genes, particularly in haploinsufficient genes and ohnologs, further underscores that the cellular abundance of the respective protein products is kept at an optimal level compatible with tightly regulated LLPS behavior, with deviations in any direction leading to serious malfunctions.

Overall, most LLPS drivers are modular proteins containing motif-rich, promiscuous, LLPS-prone IDRs of low sequence complexity, whose generally low overall abundance is under tight control. Also, their spatial distribution within cells is often highly asymmetric due to being restricted by local translation and/or anchoring mechanisms that ensure that they only reach local concentrations high enough for condensate formation at the right site(s) within cells to avoid erroneous off-site condensate formation events. In accord, one needs to take extreme care in designing and interpreting LLPS experiments, also considering that each LLPS system represents a specific case. Although measuring local protein concentrations within cells is practically out of reach, one should keep in mind that LLPS drivers might display surprisingly high local protein concentrations despite their overall low abundance. We therefore propose that proteomics-derived cellular protein concentrations should rather be treated as theoretical lower limits, not as obligatory reference points, for most *in vitro* LLPS studies.

## Author Contributions

Conceptualization, R.P., T.L., S.J.W. and P.T.; Data curation, R.P. and N.F.; Formal analysis, N.F. and R.P.; Funding acquisition, P.T.; Methodology, R.P., T.L., S.J.W. and P.T.; Resources, P.T.; Supervision, R.P., T.L., S.J.W and P.T.; Visualization, N.F.; Writing—original draft, R.P. and S.J.W.; Writing—review & editing, R.P., T.L., S.J.W. and P.T.

## References

1. Falahati, H.; Haji-Akbari, A. Thermodynamically driven assemblies and liquid–liquid phase separations in biology. Soft Matter 2019, 15, 1135–1154.

2. Alberti, S. The wisdom of crowds: regulating cell function through condensed states of living matter. J. Cell Sci. 2017, 130, 2789–2796.

3. Alberti, S.; Gladfelter, A.; Mittag, T. Considerations and Challenges in Studying Liquid-Liquid Phase Separation and Biomolecular Condensates. Cell 2019, 176, 419–434.

4. Pancsa, R.; Schad, E.; Tantos, A.; Tompa, P. Emergent functions of proteins in non-stoichiometric supramolecular assemblies. Biochim. Biophys. Acta: Proteins Proteomics 2019, doi:10.1016/j.bbapap.2019.02.007.

5. Li, X.-H.; Chavali, P.L.; Pancsa, R.; Chavali, S.; Babu, M.M. Function and Regulation of Phase-Separated Biological Condensates. Biochemistry 2018, 57, 2452–2461.

6. Banani, S.F.; Lee, H.O.; Hyman, A.A.; Rosen, M.K. Biomolecular condensates: organizers of cellular biochemistry. Nature Reviews Molecular Cell Biology 2017, 18, 285–298.

7. Al-Husini, N.; Tomares, D.T.; Bitar, O.; Childers, W.S.; Schrader, J.M. α-Proteobacterial RNA Degradosomes Assemble Liquid-Liquid Phase-Separated RNP Bodies. Mol. Cell 2018, 71, 1027–1039.e14.

8. Nikolic, J.; Le Bars, R.; Lama, Z.; Scrima, N.; Lagaudrière-Gesbert, C.; Gaudin, Y.; Blondel, D. Negri bodies are viral factories with properties of liquid organelles. Nat. Commun. 2017, 8, 58.

9. Brangwynne, C.P.; Eckmann, C.R.; Courson, D.S.; Rybarska, A.; Hoege, C.; Gharakhani, J.; Jülicher, F.; Hyman, A.A. Germline P granules are liquid droplets that localize by controlled dissolution/condensation. Science 2009, 324, 1729–1732.

10. Brangwynne, C.P.; Mitchison, T.J.; Hyman, A.A. Active liquid-like behavior of nucleoli determines their size and shape in Xenopus laevis oocytes. Proc. Natl. Acad. Sci. U. S. A. 2011, 108, 4334–4339.

11. Mitrea, D.M.; Chandra, B.; Ferrolino, M.C.; Gibbs, E.B.; Tolbert, M.; White, M.R.; Kriwacki, R.W. Methods for Physical Characterization of Phase-Separated Bodies and Membrane-less Organelles. J. Mol. Biol. 2018, 430, 4773–4805.

12. Shin, Y.; Brangwynne, C.P. Liquid phase condensation in cell physiology and disease. Science 2017, 357, doi:10.1126/science.aaf4382.

13. Alberti, S.; Dormann, D. Liquid-Liquid Phase Separation in Disease. Annu. Rev. Genet. 2019, 53, 171–194.

14. Gui, X.; Luo, F.; Li, Y.; Zhou, H.; Qin, Z.; Liu, Z.; Gu, J.; Xie, M.; Zhao, K.; Dai, B.; et al. Structural basis for reversible amyloids of hnRNPA1 elucidates their role in stress granule assembly. Nat. Commun. 2019, 10, 2006.

15. Patel, A.; Lee, H.O.; Jawerth, L.; Maharana, S.; Jahnel, M.; Hein, M.Y.; Stoynov, S.; Mahamid, J.; Saha, S.; Franzmann, T.M.; et al. A Liquid-to-Solid Phase Transition of the ALS Protein FUS Accelerated by Disease Mutation. Cell 2015, 162, 1066–1077.

16. Mann, J.R.; Gleixner, A.M.; Mauna, J.C.; Gomes, E.; DeChellis-Marks, M.R.; Needham, P.G.; Copley, K.E.; Hurtle, B.; Portz, B.; Pyles, N.J.; et al. RNA Binding Antagonizes Neurotoxic Phase Transitions of TDP-43. Neuron 2019, 102, 321–338.e8.

17. Han, T.W.; Kato, M.; Xie, S.; Wu, L.C.; Mirzaei, H.; Pei, J.; Chen, M.; Xie, Y.; Allen, J.; Xiao, G.; et al. Cell-free formation of RNA granules: bound RNAs identify features and components of cellular assemblies. Cell 2012, 149, 768–779.

18. Chen, Y.-C.M.; Kappel, C.; Beaudouin, J.; Eils, R.; Spector, D.L. Live cell dynamics of promyelocytic leukemia nuclear bodies upon entry into and exit from mitosis. Mol. Biol. Cell 2008, 19, 3147–3162.

19. Kostylev, M.A.; Tuttle, M.D.; Lee, S.; Klein, L.E.; Takahashi, H.; Cox, T.O.; Gunther, E.C.; Zilm, K.W.; Strittmatter, S.M. Liquid and Hydrogel Phases of PrP Linked to Conformation Shifts and Triggered by Alzheimer’s Amyloid-β Oligomers. Mol. Cell 2018, 72, 426–443.e12.

20. Boeynaems, S.; Alberti, S.; Fawzi, N.L.; Mittag, T.; Polymenidou, M.; Rousseau, F.; Schymkowitz, J.; Shorter, J.; Wolozin, B.; Van Den Bosch, L.; et al. Protein Phase Separation: A New Phase in Cell Biology. Trends in Cell Biology 2018, 28, 420–435.

21. Mittag, T.; Parker, R. Multiple Modes of Protein–Protein Interactions Promote RNP Granule Assembly. Journal of Molecular Biology 2018, 430, 4636–4649.

22. Roden, C.; Gladfelter, A.S. RNA contributions to the form and function of biomolecular condensates. Nat. Rev. Mol. Cell Biol. 2020, doi:10.1038/s41580-020-0264-6.

23. Langdon, E.M.; Qiu, Y.; Ghanbari Niaki, A.; McLaughlin, G.A.; Weidmann, C.A.; Gerbich, T.M.; Smith, J.A.; Crutchley, J.M.; Termini, C.M.; Weeks, K.M.; et al. mRNA structure determines specificity of a polyQ-driven phase separation. Science 2018, 360, 922–927.

24. Zhang, H.; Elbaum-Garfinkle, S.; Langdon, E.M.; Taylor, N.; Occhipinti, P.; Bridges, A.A.; Brangwynne, C.P.; Gladfelter, A.S. RNA Controls PolyQ Protein Phase Transitions. Mol. Cell 2015, 60, 220–230.

25. Hanazawa, M.; Yonetani, M.; Sugimoto, A. PGL proteins self associate and bind RNPs to mediate germ granule assembly in C. elegans. J. Cell Biol. 2011, 192, 929–937.

26. Tsang, B.; Arsenault, J.; Vernon, R.M.; Lin, H.; Sonenberg, N.; Wang, L.-Y.; Bah, A.; Forman-Kay, J.D. Phosphoregulated FMRP phase separation models activity-dependent translation through bidirectional control of mRNA granule formation. Proc. Natl. Acad. Sci. U. S. A. 2019, 116, 4218–4227.

27. Wang, J.; Choi, J.-M.; Holehouse, A.S.; Lee, H.O.; Zhang, X.; Jahnel, M.; Maharana, S.; Lemaitre, R.; Pozniakovsky, A.; Drechsel, D.; et al. A Molecular Grammar Governing the Driving Forces for Phase Separation of Prion-like RNA Binding Proteins. Cell 2018, 174, 688–699.e16.

28. Vernon, R.M.; Chong, P.A.; Tsang, B.; Kim, T.H.; Bah, A.; Farber, P.; Lin, H.; Forman-Kay, J.D. Pi-Pi contacts are an overlooked protein feature relevant to phase separation. Elife 2018, 7, doi:10.7554/eLife.31486.

29. Bratek-Skicki, A.; Pancsa, R.; Meszaros, B.; Van Lindt, J.; Tompa, P. A guide to regulation of the formation of biomolecular condensates. FEBS J. 2020, 287, 1924–1935.

30. Yoo, H.; Triandafillou, C.; Drummond, D.A. Cellular sensing by phase separation: Using the process, not just the products. J. Biol. Chem. 2019, 294, 7151–7159.

31. Söding, J.; Zwicker, D.; Sohrabi-Jahromi, S.; Boehning, M.; Kirschbaum, J. Mechanisms for Active Regulation of Biomolecular Condensates. Trends Cell Biol. 2020, 30, 4–14.

32. Owen, I.; Shewmaker, F. The Role of Post-Translational Modifications in the Phase Transitions of Intrinsically Disordered Proteins. Int. J. Mol. Sci. 2019, 20, doi:10.3390/ijms20215501.

33. Snead, W.T.; Gladfelter, A.S. The Control Centers of Biomolecular Phase Separation: How Membrane Surfaces, PTMs, and Active Processes Regulate Condensation. Mol. Cell 2019, 76, 295–305.

34. Willadsen, K.; Mohamad, N.; Bodén, M. NSort/DB: an intranuclear compartment protein database. Genomics Proteomics Bioinformatics 2012, 10, 226–229.

35. Nunes, C.; Mestre, I.; Marcelo, A.; Koppenol, R.; Matos, C.A.; Nóbrega, C. MSGP: the first database of the protein components of the mammalian stress granules. Database 2019, 2019, doi:10.1093/database/baz031.

36. Youn, J.-Y.; Dyakov, B.J.A.; Zhang, J.; Knight, J.D.R.; Vernon, R.M.; Forman-Kay, J.D.; Gingras, A.-C. Properties of Stress Granule and P-Body Proteomes. Mol. Cell 2019, 76, 286–294.

37. Mészáros, B.; Erdős, G.; Szabó, B.; Schád, É.; Tantos, Á.; Abukhairan, R.; Horváth, T.; Murvai, N.; Kovács, O.P.; Kovács, M.; et al. PhaSePro: the database of proteins driving liquid-liquid phase separation. Nucleic Acids Res. 2020, 48, D360–D367.

38. Li, Q.; Peng, X.; Li, Y.; Tang, W.; Zhu, J. ’an; Huang, J.; Qi, Y.; Zhang, Z. LLPSDB: a database of proteins undergoing liquid-liquid phase separation in vitro. Nucleic Acids Res. 2020, 48, D320–D327.

39. Ning, W.; Guo, Y.; Lin, S.; Mei, B.; Wu, Y.; Jiang, P.; Tan, X.; Zhang, W.; Chen, G.; Peng, D.; et al. DrLLPS: a data resource of liquid-liquid phase separation in eukaryotes. Nucleic Acids Res. 2020, 48, D288–D295.

40. You, K.; Huang, Q.; Yu, C.; Shen, B.; Sevilla, C.; Shi, M.; Hermjakob, H.; Chen, Y.; Li, T. PhaSepDB: a database of liquid-liquid phase separation related proteins. Nucleic Acids Res. 2020, 48, D354–D359.

41. Li, Q.; Wang, X.; Dou, Z.; Yang, W.; Huang, B.; Lou, J.; Zhang, Z. Protein Databases Related to Liquid-Liquid Phase Separation. Int. J. Mol. Sci. 2020, 21, doi:10.3390/ijms21186796.

42. Pancsa, R.; Vranken, W.; Mészáros, B. Computational resources for identifying and describing proteins driving liquid-liquid phase separation. Brief. Bioinform. 2021, doi:10.1093/bib/bbaa408.

43. Turinsky, A.L.; Razick, S.; Turner, B.; Donaldson, I.M.; Wodak, S.J. Literature curation of protein interactions: measuring agreement across major public databases. Database 2010, 2010, baq026.

44. Turinsky, A.L.; Razick, S.; Turner, B.; Donaldson, I.M.; Wodak, S.J. Interaction databases on the same page. Nat. Biotechnol. 2011, 29, 391–393.

45. Su, X.; Ditlev, J.A.; Hui, E.; Xing, W.; Banjade, S.; Okrut, J.; King, D.S.; Taunton, J.; Rosen, M.K.; Vale, R.D. Phase separation of signaling molecules promotes T cell receptor signal transduction. Science 2016, 352, 595–599.

46. Mitrea, D.M.; Cika, J.A.; Guy, C.S.; Ban, D.; Banerjee, P.R.; Stanley, C.B.; Nourse, A.; Deniz, A.A.; Kriwacki, R.W. Nucleophosmin integrates within the nucleolus via multi-modal interactions with proteins displaying R-rich linear motifs and rRNA. Elife 2016, 5, doi:10.7554/eLife.13571.

47. Bouchard, J.J.; Otero, J.H.; Scott, D.C.; Szulc, E.; Martin, E.W.; Sabri, N.; Granata, D.; Marzahn, M.R.; Lindorff-Larsen, K.; Salvatella, X.; et al. Cancer Mutations of the Tumor Suppressor SPOP Disrupt the Formation of Active, Phase-Separated Compartments. Mol. Cell 2018, 72, 19–36.e8.

48. Banani, S.F.; Rice, A.M.; Peeples, W.B.; Lin, Y.; Jain, S.; Parker, R.; Rosen, M.K. Compositional Control of Phase-Separated Cellular Bodies. Cell 2016, 166, 651–663.

49. Bolognesi, B.; Lorenzo Gotor, N.; Dhar, R.; Cirillo, D.; Baldrighi, M.; Tartaglia, G.G.; Lehner, B. A Concentration-Dependent Liquid Phase Separation Can Cause Toxicity upon Increased Protein Expression. Cell Rep. 2016, 16, 222–231.

50. Cinar, S.; Cinar, H.; Chan, H.S.; Winter, R. Pressure-Sensitive and Osmolyte-Modulated Liquid-Liquid Phase Separation of Eye-Lens γ-Crystallins. J. Am. Chem. Soc. 2019, 141, 7347–7354.

51. Kato, M.; Han, T.W.; Xie, S.; Shi, K.; Du, X.; Wu, L.C.; Mirzaei, H.; Goldsmith, E.J.; Longgood, J.; Pei, J.; et al. Cell-free formation of RNA granules: low complexity sequence domains form dynamic fibers within hydrogels. Cell 2012, 149, 753–767.

52. Boija, A.; Klein, I.A.; Sabari, B.R.; Dall’Agnese, A.; Coffey, E.L.; Zamudio, A.V.; Li, C.H.; Shrinivas, K.; Manteiga, J.C.; Hannett, N.M.; et al. Transcription Factors Activate Genes through the Phase-Separation Capacity of Their Activation Domains. Cell 2018, 175, 1842–1855.e16.

53. Ying, Y.; Wang, X.-J.; Vuong, C.K.; Lin, C.-H.; Damianov, A.; Black, D.L. Splicing Activation by Rbfox Requires Self-Aggregation through Its Tyrosine-Rich Domain. Cell 2017, 170, 312–323.e10.

54. Lin, Y.; Protter, D.S.W.; Rosen, M.K.; Parker, R. Formation and Maturation of Phase-Separated Liquid Droplets by RNA-Binding Proteins. Mol. Cell 2015, 60, 208–219.

55. Wang, M.; Herrmann, C.J.; Simonovic, M.; Szklarczyk, D.; von Mering, C. Version 4.0 of PaxDb: Protein abundance data, integrated across model organisms, tissues, and cell-lines. Proteomics 2015, 15, 3163–3168.

56. Milo, R. What is the total number of protein molecules per cell volume? A call to rethink some published values. Bioessays 2013, 35, 1050–1055.

57. Hennig, S.; Kong, G.; Mannen, T.; Sadowska, A.; Kobelke, S.; Blythe, A.; Knott, G.J.; Iyer, K.S.; Ho, D.; Newcombe, E.A.; et al. Prion-like domains in RNA binding proteins are essential for building subnuclear paraspeckles. J. Cell Biol. 2015, 210, 529–539.

58. Milles, S.; Lemke, E.A. Single molecule study of the intrinsically disordered FG-repeat nucleoporin 153. Biophys. J. 2011, 101, 1710–1719.

59. Lu, H.; Yu, D.; Hansen, A.S.; Ganguly, S.; Liu, R.; Heckert, A.; Darzacq, X.; Zhou, Q. Phase-separation mechanism for C-terminal hyperphosphorylation of RNA polymerase II. Nature 2018, 558, 318–323.

60. Lee, H.Y.; Kim, E.G.; Jung, H.R.; Jung, J.W.; Kim, H.B.; Cho, J.W.; Kim, K.M.; Yi, E.C. Refinements of LC-MS/MS Spectral Counting Statistics Improve Quantification of Low Abundance Proteins. Sci. Rep. 2019, 9, 13653.

61. Lundgren, D.H.; Hwang, S.-I.; Wu, L.; Han, D.K. Role of spectral counting in quantitative proteomics. Expert Rev. Proteomics 2010, 7, 39–53.

62. Smaczniak, C.; Li, N.; Boeren, S.; America, T.; van Dongen, W.; Goerdayal, S.S.; de Vries, S.; Angenent, G.C.; Kaufmann, K. Proteomics-based identification of low-abundance signaling and regulatory protein complexes in native plant tissues. Nat. Protoc. 2012, 7, 2144–2158.

63. Stergachis, A.B.; MacLean, B.; Lee, K.; Stamatoyannopoulos, J.A.; MacCoss, M.J. Rapid empirical discovery of optimal peptides for targeted proteomics. Nat. Methods 2011, 8, 1041–1043.

64. Alfonso-Garrido, J.; Garcia-Calvo, E.; Luque-Garcia, J.L. Sample preparation strategies for improving the identification of membrane proteins by mass spectrometry. Anal. Bioanal. Chem. 2015, 407, 4893–4905.

65. Zeng, M.; Bai, G.; Zhang, M. Anchoring high concentrations of SynGAP at postsynaptic densities via liquid-liquid phase separation. Small GTPases 2019, 10, 296–304.

66. Ding, C.; Chan, D.W.; Liu, W.; Liu, M.; Li, D.; Song, L.; Li, C.; Jin, J.; Malovannaya, A.; Jung, S.Y.; et al. Proteome-wide profiling of activated transcription factors with a concatenated tandem array of transcription factor response elements. Proc. Natl. Acad. Sci. U. S. A. 2013, 110, 6771–6776.

67. Wierer, M.; Mann, M. Proteomics to study DNA-bound and chromatin-associated gene regulatory complexes. Hum. Mol. Genet. 2016, 25, R106–R114.

68. Trinidad, J.C.; Thalhammer, A.; Specht, C.G.; Lynn, A.J.; Baker, P.R.; Schoepfer, R.; Burlingame, A.L. Quantitative analysis of synaptic phosphorylation and protein expression. Mol. Cell. Proteomics 2008, 7, 684–696.

69. Collins, M.O.; Yu, L.; Coba, M.P.; Husi, H.; Campuzano, I.; Blackstock, W.P.; Choudhary, J.S.; Grant, S.G.N. Proteomic analysis of in vivo phosphorylated synaptic proteins. J. Biol. Chem. 2005, 280, 5972–5982.

70. Levy, E.D.; Kowarzyk, J.; Michnick, S.W. High-resolution mapping of protein concentration reveals principles of proteome architecture and adaptation. Cell Rep. 2014, 7, 1333–1340.

71. Boisvert, F.-M.; Lam, Y.W.; Lamont, D.; Lamond, A.I. A quantitative proteomics analysis of subcellular proteome localization and changes induced by DNA damage. Mol. Cell. Proteomics 2010, 9, 457–470.

72. Gatto, L.; Breckels, L.M.; Lilley, K.S. Assessing sub-cellular resolution in spatial proteomics experiments. Curr. Opin. Chem. Biol. 2019, 48, 123–149.

73. Holt, C.E.; Martin, K.C.; Schuman, E.M. Local translation in neurons: visualization and function. Nat. Struct. Mol. Biol. 2019, 26, 557–566.

74. Cajigas, I.J.; Tushev, G.; Will, T.J.; tom Dieck, S.; Fuerst, N.; Schuman, E.M. The local transcriptome in the synaptic neuropil revealed by deep sequencing and high-resolution imaging. Neuron 2012, 74, 453–466.

75. Wang, T.; Hamilla, S.; Cam, M.; Aranda-Espinoza, H.; Mili, S. Extracellular matrix stiffness and cell contractility control RNA localization to promote cell migration. Nat. Commun. 2017, 8, 896.

76. Weatheritt, R.J.; Gibson, T.J.; Babu, M.M. Asymmetric mRNA localization contributes to fidelity and sensitivity of spatially localized systems. Nat. Struct. Mol. Biol. 2014, 21, 833–839.

77. Wilhelm, B.G.; Mandad, S.; Truckenbrodt, S.; Kröhnert, K.; Schäfer, C.; Rammner, B.; Koo, S.J.; Claßen, G.A.; Krauss, M.; Haucke, V.; et al. Composition of isolated synaptic boutons reveals the amounts of vesicle trafficking proteins. Science 2014, 344, 1023–1028.

78. Sugiyama, Y.; Kawabata, I.; Sobue, K.; Okabe, S. Determination of absolute protein numbers in single synapses by a GFP-based calibration technique. Nat. Methods 2005, 2, 677–684.

79. Zeng, M.; Shang, Y.; Araki, Y.; Guo, T.; Huganir, R.L.; Zhang, M. Phase Transition in Postsynaptic Densities Underlies Formation of Synaptic Complexes and Synaptic Plasticity. Cell 2016, 166, 1163–1175.e12.

80. Milovanovic, D.; Wu, Y.; Bian, X.; De Camilli, P. A liquid phase of synapsin and lipid vesicles. Science 2018, 361, 604–607.

81. Tari, M.; Manceau, V.; de Matha Salone, J.; Kobayashi, A.; Pastré, D.; Maucuer, A. U2AF assemblies drive sequence-specific splice site recognition. EMBO Rep. 2019, 20, e47604.

82. Ries, R.J.; Zaccara, S.; Klein, P.; Olarerin-George, A.; Namkoong, S.; Pickering, B.F.; Patil, D.P.; Kwak, H.; Lee, J.H.; Jaffrey, S.R. mA enhances the phase separation potential of mRNA. Nature 2019, 571, 424–428.

83. Wang, M.; Tao, X.; Jacob, M.D.; Bennett, C.A.; Ho, J.J.D.; Gonzalgo, M.L.; Audas, T.E.; Lee, S. Stress-Induced Low Complexity RNA Activates Physiological Amyloidogenesis. Cell Rep. 2018, 24, 1713–1721.e4.

84. Boehning, M.; Dugast-Darzacq, C.; Rankovic, M.; Hansen, A.S.; Yu, T.; Marie-Nelly, H.; McSwiggen, D.T.; Kokic, G.; Dailey, G.M.; Cramer, P.; et al. RNA polymerase II clustering through carboxy-terminal domain phase separation. Nat. Struct. Mol. Biol. 2018, 25, 833–840.

85. Zhang, Y.; Bertulat, B.; Tencer, A.H.; Ren, X.; Wright, G.M.; Black, J.; Cardoso, M.C.; Kutateladze, T.G. MORC3 Forms Nuclear Condensates through Phase Separation. iScience 2019, 17, 182–189.

86. Nair, S.J.; Yang, L.; Meluzzi, D.; Oh, S.; Yang, F.; Friedman, M.J.; Wang, S.; Suter, T.; Alshareedah, I.; Gamliel, A.; et al. Phase separation of ligand-activated enhancers licenses cooperative chromosomal enhancer assembly. Nat. Struct. Mol. Biol. 2019, 26, 193–203.

87. Schmidt, H.B.; Görlich, D. Nup98 FG domains from diverse species spontaneously phase-separate into particles with nuclear pore-like permselectivity. Elife 2015, 4, doi:10.7554/eLife.04251.

88. Larson, A.G.; Elnatan, D.; Keenen, M.M.; Trnka, M.J.; Johnston, J.B.; Burlingame, A.L.; Agard, D.A.; Redding, S.; Narlikar, G.J. Liquid droplet formation by HP1α suggests a role for phase separation in heterochromatin. Nature 2017, 547, 236–240.

89. Sabari, B.R.; Dall’Agnese, A.; Boija, A.; Klein, I.A.; Coffey, E.L.; Shrinivas, K.; Abraham, B.J.; Hannett, N.M.; Zamudio, A.V.; Manteiga, J.C.; et al. Coactivator condensation at super-enhancers links phase separation and gene control. Science 2018, 361, doi:10.1126/science.aar3958.

90. Shrinivas, K.; Sabari, B.R.; Coffey, E.L.; Klein, I.A.; Boija, A.; Zamudio, A.V.; Schuijers, J.; Hannett, N.M.; Sharp, P.A.; Young, R.A.; et al. Enhancer Features that Drive Formation of Transcriptional Condensates. Mol. Cell 2019, 75, 549–561.e7.

91. Strom, A.R.; Emelyanov, A.V.; Mir, M.; Fyodorov, D.V.; Darzacq, X.; Karpen, G.H. Phase separation drives heterochromatin domain formation. Nature 2017, 547, 241–245.

92. Wang, L.; Gao, Y.; Zheng, X.; Liu, C.; Dong, S.; Li, R.; Zhang, G.; Wei, Y.; Qu, H.; Li, Y.; et al. Histone Modifications Regulate Chromatin Compartmentalization by Contributing to a Phase Separation Mechanism. Mol. Cell 2019, 76, 646–659.e6.

93. Molliex, A.; Temirov, J.; Lee, J.; Coughlin, M.; Kanagaraj, A.P.; Kim, H.J.; Mittag, T.; Taylor, J.P. Phase separation by low complexity domains promotes stress granule assembly and drives pathological fibrillization. Cell 2015, 163, 123–133.

94. Kaur, T.; Alshareedah, I.; Wang, W.; Ngo, J.; Moosa, M.M.; Banerjee, P.R. Molecular Crowding Tunes Material States of Ribonucleoprotein Condensates. Biomolecules 2019, 9, doi:10.3390/biom9020071.

95. Walter, H.; Brooks, D.E. Phase separation in cytoplasm, due to macromolecular crowding, is the basis for microcompartmentation. FEBS Lett. 1995, 361, 135–139.

96. Protter, D.S.W.; Rao, B.S.; Van Treeck, B.; Lin, Y.; Mizoue, L.; Rosen, M.K.; Parker, R. Intrinsically Disordered Regions Can Contribute Promiscuous Interactions to RNP Granule Assembly. Cell Rep. 2018, 22, 1401–1412.

97. McSwiggen, D.T.; Mir, M.; Darzacq, X.; Tjian, R. Evaluating phase separation in live cells:diagnosis, caveats, and functional consequences. Genes Dev. 2019, 33, 1619–1634.

98. Schütz, S.; Nöldeke, E.R.; Sprangers, R. A synergistic network of interactions promotes the formation of in vitro processing bodies and protects mRNA against decapping. Nucleic Acids Res. 2017, 45, 6911–6922.

99. Banjade, S.; Wu, Q.; Mittal, A.; Peeples, W.B.; Pappu, R.V.; Rosen, M.K. Conserved interdomain linker promotes phase separation of the multivalent adaptor protein Nck. Proc. Natl. Acad. Sci. U. S. A. 2015, 112, E6426–35.

100. Li, P.; Banjade, S.; Cheng, H.-C.; Kim, S.; Chen, B.; Guo, L.; Llaguno, M.; Hollingsworth, J.V.; King, D.S.; Banani, S.F.; et al. Phase transitions in the assembly of multivalent signalling proteins. Nature 2012, 483, 336–340.

101. Gsponer, J.; Futschik, M.E.; Teichmann, S.A.; Babu, M.M. Tight Regulation of Unstructured Proteins: From Transcript Synthesis to Protein Degradation. Science 2008, 322, 1365–1368.

102. Vavouri, T.; Semple, J.I.; Garcia-Verdugo, R.; Lehner, B. Intrinsic protein disorder and interaction promiscuity are widely associated with dosage sensitivity. Cell 2009, 138, 198–208.

103. Ni, Z.; Zhou, X.-Y.; Aslam, S.; Niu, D.-K. Characterization of Human Dosage-Sensitive Transcription Factor Genes. Front. Genet. 2019, 10, 1208.

104. The UniProt Consortium UniProt: the universal protein knowledgebase. Nucleic Acids Res. 2017, 45, D158–D169.

105. Lek, M.; Karczewski, K.J.; Minikel, E.V.; Samocha, K.E.; Banks, E.; Fennell, T.; O’Donnell-Luria, A.H.; Ware, J.S.; Hill, A.J.; Cummings, B.B.; et al. Analysis of protein-coding genetic variation in 60,706 humans. Nature 2016, 536, 285–291.

106. Shihab, H.A.; Rogers, M.F.; Campbell, C.; Gaunt, T.R. HIPred: an integrative approach to predicting haploinsufficient genes. Bioinformatics 2017, 33, 1751–1757.

107. Makino, T.; McLysaght, A. Ohnologs in the human genome are dosage balanced and frequently associated with disease. Proc. Natl. Acad. Sci. U. S. A. 2010, 107, 9270–9274.

108. Rice, A.M.; McLysaght, A. Dosage sensitivity is a major determinant of human copy number variant pathogenicity. Nat. Commun. 2017, 8, 14366.

109. Lambert, S.A.; Jolma, A.; Campitelli, L.F.; Das, P.K.; Yin, Y.; Albu, M.; Chen, X.; Taipale, J.; Hughes, T.R.; Weirauch, M.T. The Human Transcription Factors. Cell 2018, 175, 598–599.

110. Gutierrez, M.G.; Malmstadt, N. Human serotonin receptor 5-HT(1A) preferentially segregates to the liquid disordered phase in synthetic lipid bilayers. J. Am. Chem. Soc. 2014, 136, 13530–13533.

111. Dubreuil, B.; Matalon, O.; Levy, E.D. Protein Abundance Biases the Amino Acid Composition of Disordered Regions to Minimize Non-functional Interactions. J. Mol. Biol. 2019, 431, 4978–4992.

112. Wiśniewski, J.R.; Hein, M.Y.; Cox, J.; Mann, M. A “proteomic ruler” for protein copy number and concentration estimation without spike-in standards. Mol. Cell. Proteomics 2014, 13, 3497–3506.

