## Supplementary Figures for "Concentration and dosage sensitivity of proteins driving liquid-liquid phase separation"

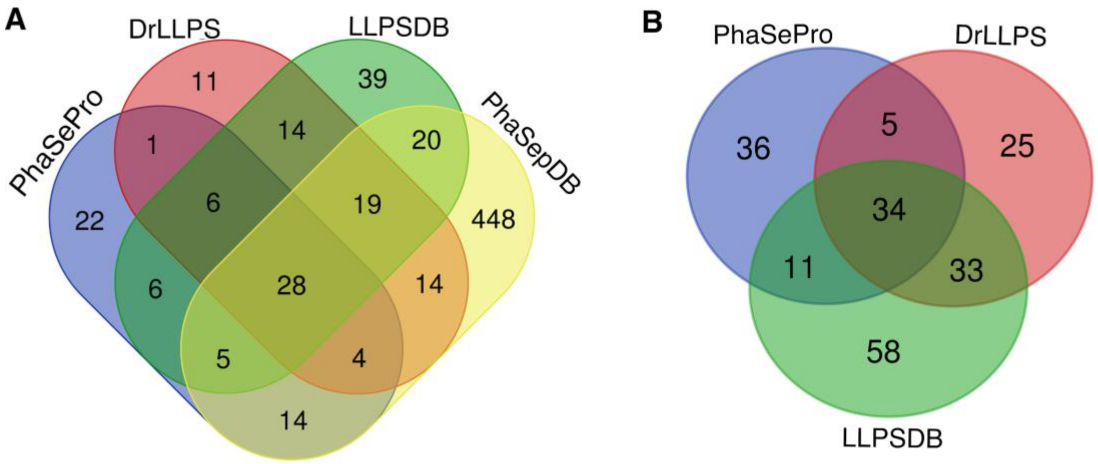


**Supplementary Figure S1: Overlap of the literature references addressing the investigated entries.** A) The Venn diagram compares the literature references supporting the localization to MLOs/ ability to undergo LLPS of the selected subsets of LLPS-associated proteins across the four resources. B) The same Venn diagram is shown for the three resources that could be analyzed further.

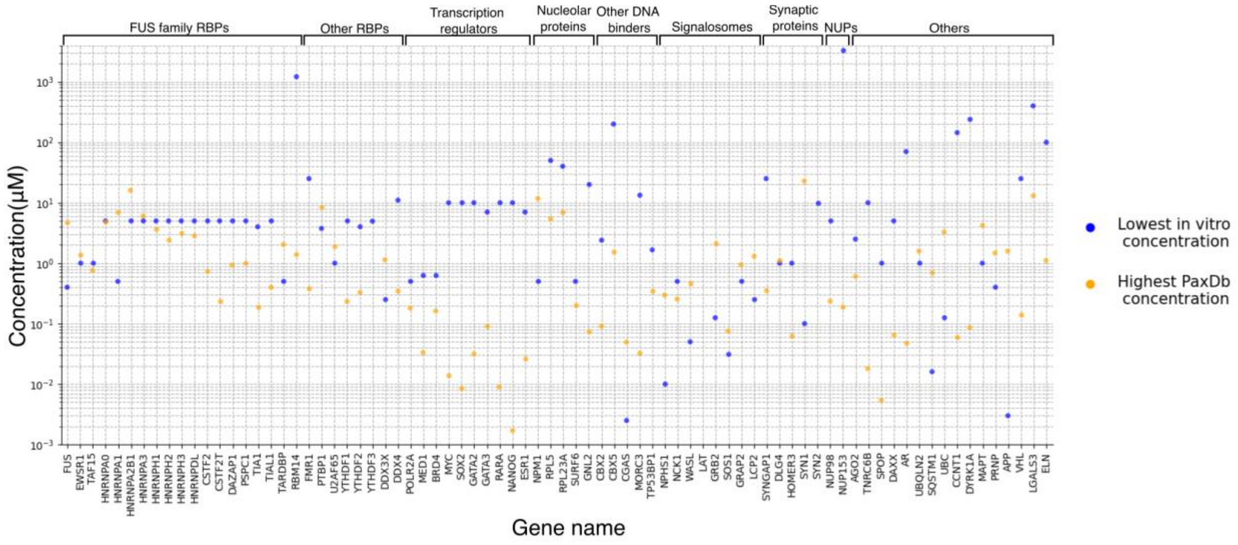


**Supplementary Figure S2: Lowest protein concentrations applied in *in vitro* LLPS experiments frequently exceed those calculated from proteomics-derived highest cellular protein abundances.** Comparison of the lowest protein concentrations applied in *in vitro* LLPS experiments where the given proteins could phase separate (in blue) to the respective cellular concentrations calculated from proteomics-derived highest protein abundances stored in PaxDb (in orange) for the 78 true human LLPS (co)driver proteins backed by *in vitro* experiments. Only integrated datasets were considered for PaxDb. Concentrations are provided in micromolar units, while LLPS proteins are represented by the respective gene names on the horizontal axis. Functional groups of the proteins are indicated on the top of the figure.

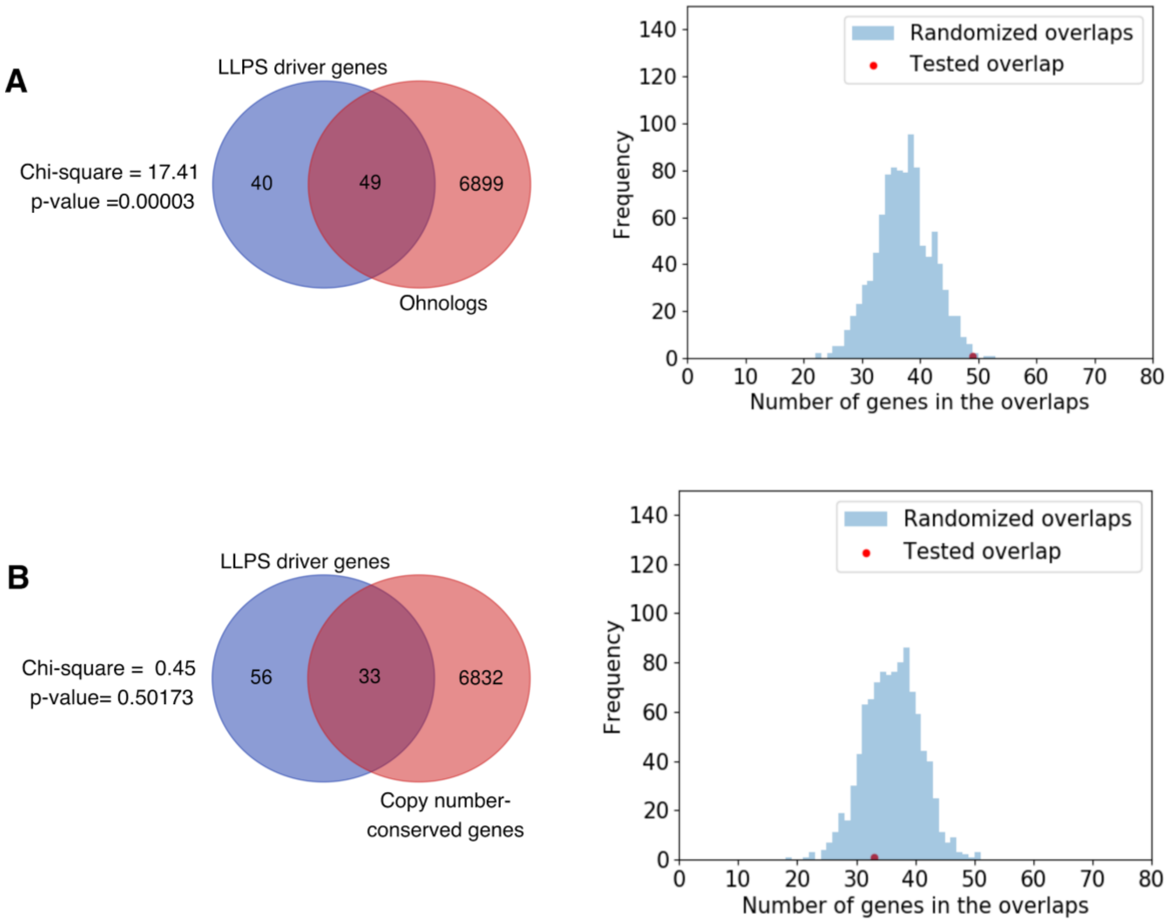


**Supplementary Figure S3: The genes of human LLPS driver proteins are overrepresented among ohnologs but not among copy number-conserved genes.** Enrichment analysis of human LLPS driver genes among (A) ohnologs and (B) copy number-conserved genes. The Venn diagrams show the overlap between human LLPS driver genes and (A) ohnologs or (B) copy number-conserved genes. The detected overlap is only significant for ohnologs based on chi$^2$ tests using the whole human reviewed UniProt proteome as background (left). Histograms showing the difference between the overlaps of human LLPS driver genes (red) or equivalent sets of randomly selected well-annotated genes (blue) with (A) ohnologs or (B) copy number-conserved genes.

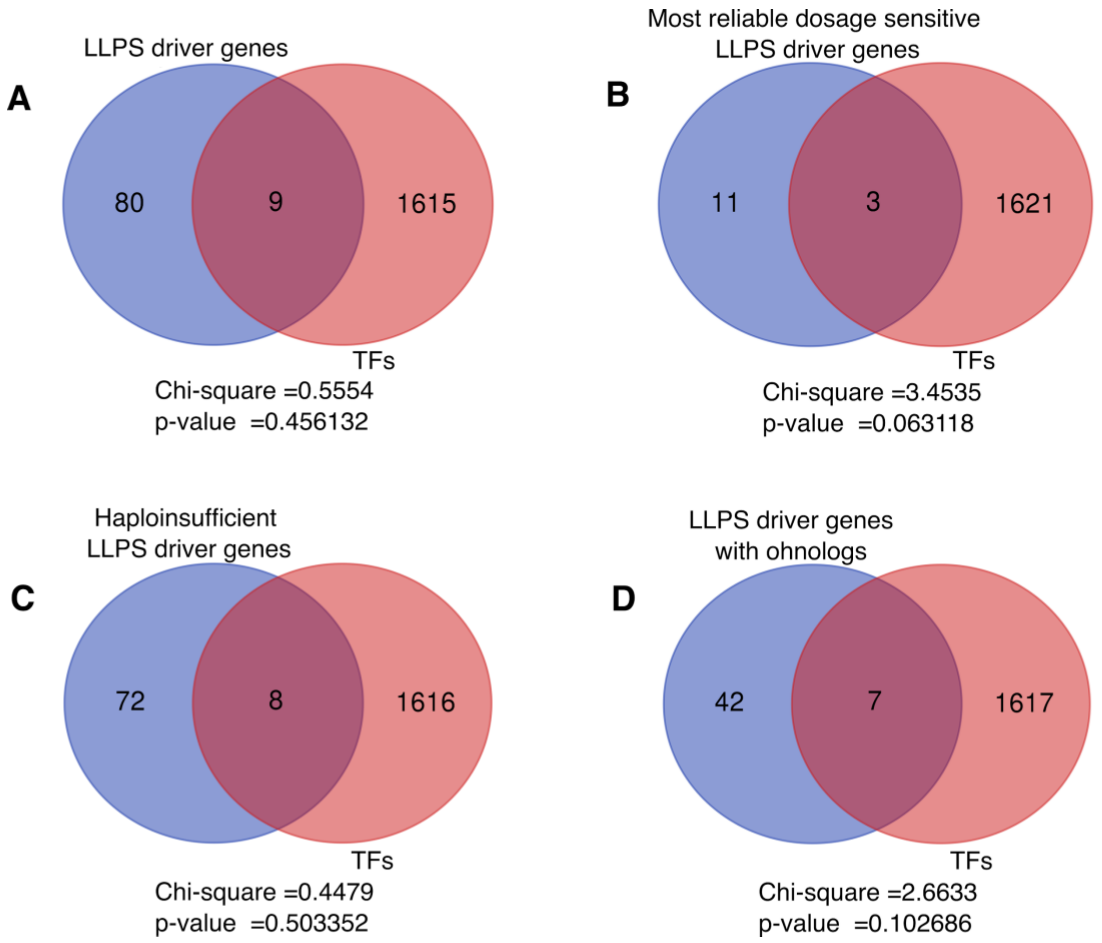


**Supplementary Figure S4: Lack of enrichment in transcription factors among LLPS driver genes.** The Venn diagrams show the overlap between the transcription factor (TF) genes and the human LLPS driver genes (A), the set of most reliable dosage sensitive (MRDS) LLPS driver genes (B), the set of haploinsufficient LLPS driver genes (C), and the set of LLPS driver genes with ohnologs (D). The detected overlaps do not represent statistically significant enrichments based on the $chi^2$ tests.
